# Transcriptomic analysis of FER-RALF-LRX pathway mutants suggests constitutive gene expression defects contribute to powdery mildew resistance

**DOI:** 10.64898/2026.06.25.734470

**Authors:** Henriette Leicher, Amit Fenn, Maxim Messerer, Christine Wurmser, Ralph Hückelhoven, Nadia Kamal, Martin Stegmann

## Abstract

The receptor kinase FERONIA (FER) perceives endogenous RAPID ALKALINIZATION FACTOR (RALF) peptides and regulates a plethora of plant physiological processes, including immunity. RALF peptides also bind to LEUCINE-RICH REPEAT EXTENSIN (LRX) proteins as structural components of the cell wall. We recently showed that the FER-RALF-LRX pathway supports colonization by the obligate biotrophic fungal pathogen *Erysiphe cruciferarum (Ecr)*, a member of the powdery mildew species complex that infects Arabidopsis. Genetic disruption of the pathway primarily affects conidiation of the fungus, raising the question of effects on fungal nutrition. To get further insight into the underlying mechanisms, we performed RNA sequencing (RNAseq) to identify differential transcriptional responses of FER-RALF-LRX pathway mutants upon *Ecr* infection. Surprisingly, our results revealed that pathway disruption has a limited impact on the overall transcriptional changes upon fungal infection. However, consistent with previous reports, FER-RALF-LRX pathway mutants show changes in basal expression of a plethora of genes, mainly associated with cell wall metabolism, jasmonic acid signalling, amino acid biosynthesis and secondary metabolism. Many of these genes are regulated by *Ecr* infection across genotypes, too. This raises the question whether these are relevant pathway components for powdery mildew host establishment downstream of the FER-RALF-LRX module. In summary, our data reveals new insights into FER-RALF-LRX-dependent responses that may support host susceptibility to biotrophic plant pathogens.

## Introduction

Powdery mildew fungi are obligate biotrophic ascomycetes that belong to the order Erysiphales. They infect a wide range of plant species, including many economically important crops. Infection leads to significant losses in yield quantity and quality (Glawe, 2008). Because of their strict biotrophic lifestyle, powdery mildew fungi rely on nutrient uptake from living host cells, which requires a high degree of adaptation to the host plant (Eichmann & Hückelhoven, 2008). This has led to the evolution of highly host-specialized powdery mildew species and *formae speciales*. *Blumeria hordei* (*Bh*) is virulent on barley but fails to infect *Arabidopsis thaliana* (hereafter Arabidopsis). *Golovinomyces cichoracearum* (syn. *Erysiphe cichoracearum*), *G. orontii* and *E. cruciferarum* are three examples of powdery mildew fungi that infect Arabidopsis (Lipka *et al*., 2008). Host specialisation suggests the necessity for a high degree of adaptation to achieve molecular manipulation of host physiology and successful colonisation.

Powdery mildew infection starts with a conidiospore landing on the leaf surface, followed by spore germination, the formation of a germ tube and appressorium. These structures are required to apply pressure and secret enzymes for cuticle and cell wall penetration (Eichmann & Hückelhoven, 2008; Glawe, 2008). The plant can stop this infection attempt through the formation of papillae, which reinforce cell walls by localised deposition of callose and other polysaccharides, as well as cross-linking of cell wall material at the site of penetration attempt (Chowdhury *et al*., 2014). Besides structural reinforcement of the cell wall, tryptophan-derived secondary metabolites appear crucial to limit powdery mildew invasion of Arabidopsis (Consonni *et al*., 2009; Hunziker *et al*., 2020). Successful penetration is followed by the formation of a haustorium. This specialised hyphae invaginates the host plasma membrane without disrupting its integrity (Eichmann & Hückelhoven, 2008). Consequently, the haustorium is surrounded by an extrahaustorial membrane (EHM), which is plant-derived and has distinct protein and lipid compositions (Hückelhoven & Panstruga, 2011). Haustoria formation is accompanied by extensive transcriptional changes in the infected tissue, as well as reorganisation of the actin cytoskeleton and the endomembrane system (Micali *et al*., 2008; Hückelhoven & Panstruga, 2011). Once the haustorium is established and the fungus can obtain sufficient nutrients, it continues to grow epiphytically by elongating secondary hyphae that penetrate neighbouring cells. In the final stage of infection, conidiophores emerge from the epiphytic mycelium and produce new conidiospores, completing powdery mildew’s asexual life cycle (Eichmann & Hückelhoven, 2008). As newly produced conidiospores store energy for the next infection cycle, it is thought that this stage of infection requires high levels of nutrient uptake from the host (Zeng *et al*., 2017).

The successful establishment of a biotrophic interaction depends on pathogen-derived effectors that modulate host physiology and host susceptibility factors that support fungal host establishment and growth (Micali *et al*., 2008). Susceptibility factor loss-of-function mutants are more resistant to infection. The most prominent example is MILDEW RESISTANCE LOCUS O (MLO) with mutants displaying durable and broad-spectrum penetration resistance to powdery mildew across plant lineages (Kusch & Panstruga, 2017). Other susceptibility factors, such as POWDERY MILDEW RESISTANT 5 (PMR5) and PMR6, do not strongly affect host penetration but limit fungal growth and conidiophore production. The *pmr5* and *pmr6* mutants do not show enhanced immune responses and their resistance is largely independent of the classical defense hormones salicylic acid (SA), jasmonic acid (JA) and ethylene. Instead, they exhibit altered cell wall composition, which may reduce their suitability as a host (Vogel *et al*., 2002; Vogel *et al*., 2004; Chiniquy *et al*., 2019). The mechanistic basis of this, however, remains unknown.

An important example of a susceptibility factor in plant powdery mildew interaction that affects post-invasive fungal growth without controlling host penetration is the receptor-like kinase FERONIA (FER) (Kessler *et al*., 2010; Leicher *et al*., 2026). Loss of *FER* results in a reduction of powdery mildew conidiophore production without obvious alterations in SA signalling (Leicher *et al*., 2026). Notably, *fer*-4 mutants show enhanced susceptibility to infection with *Pseudomonas syringae* pv. *tomato* DC3000 (*Pto*) and *Hyaloperonospora arabidopsidis (Stegmann et al., 2017; Leicher et al., 2026)*, suggesting a powdery mildew-specific susceptibility function. FER perceives endogenous RAPID ALKALINIZATION FACTOR (RALF) peptides. RALF peptides can also bind de-methylated cell wall pectins via cell wall-localised LEUCINE-RICH REPEAT (LRX), revealing a dual function as cell wall structural components and signalling molecules with both functions being tightly interconnected (Moussu *et al*., 2023; Chen *et al*., 2024; Rößling *et al*., 2024; Schoenaers *et al*., 2024; Biermann *et al*., 2025; Schade *et al*., 2025). Importantly, RALF peptides and LRX proteins also act as susceptibility factors for powdery mildew infection (Leicher *et al*., 2026).

The FER-RALF-LRX-signalling module is involved in multiple physiological processes, including cell expansion, reproduction and stress responses (Zhang *et al*., 2020). Further, FER is part of the cell wall integrity surveillance machinery (Chen *et al*., 2024; Cheung, 2024). Consistent with this, fer-4 mutants exhibit altered cell wall compositions with reduced cellulose content (Yeats *et al*., 2016; Biermann *et al*., 2025). Moreover, FER represses JA-signalling by destabilising the JA-regulated transcription factor MYC2 (Guo *et al*., 2018). In addition to enhanced JA-signalling in *fer-4* mutants, a recent study showed that genes involved in indole glucosinolate biosynthesis are upregulated in *fer-4* (Wang *et al*., 2022). Interestingly, these secondary metabolites are linked to non-host resistance against powdery mildew (Bednarek *et al*., 2009; Hunziker *et al*., 2020).

The complexity of FER-RALF-LRX-modulated responses poses a challenge in pinpointing specific pathways that support powdery mildew infection. In a recent study, we demonstrated that apoplastic pH modulation and cell wall remodelling, two key features of FER/RALF-signalling, regulate powdery mildew conidiation (Leicher *et al*., 2026). By investigating gene expression via RNA sequencing (RNAseq), we now aim to gain a better understanding of the FER-RALF-LRX-dependent transcriptional response during powdery mildew infection. We performed gene expression analysis at 1 and 4 days post-infection (dpi) with the powdery mildew fungus *Erysiphe cruciferarum* on *fer-4*, *lrx1/2/3/4/5 (lrx5x*) and *ralf7x* (CRISPR-induced mutations in *RALF1, RALF18, RALF22, RALF23, RALF31, RALF33, RALF34*). Analysing FER-RALF-LRX signalling mutants during biotrophic fungal reproduction provides an opportunity to investigate factors controlling post-penetration colonization in a compatible interaction.

Our data shows that the transcriptional response to powdery mildew infection is similar between wildtype and *fer-4/lrx5x*/*ralf7x*. All genotypes show comparable upregulation of defense genes and downregulation of photosynthesis-related genes. The similar transcriptomic profile upon infection raised the question whether steady-state gene expression differences between wild type and mutant explains their powdery mildew resistance phenotype. Basal differences in the transcriptome of FER-RALF-LRX-mutant plants revealed an upregulation of JA-signalling and JA-dependent secondary metabolism as well as a downregulation of cell wall metabolic processes. Interestingly, many of these genes are regulated in the wild type upon powdery mildew infection, too. Therefore, we hypothesize that altered metabolism and cell wall composition underlines the resistance phenotype of RALF pathway mutants.

## Material and Methods

### Powdery mildew propagation

*Erysiphe cruciferarum* was maintained by weekly infection of 4 – 5-week-old uninfected Col-0 and *pad4* plants. Spores were used for further propagation 3 – 4 weeks after inoculation. Powdery mildew propagation and infection experiments were performed in an environmentally controlled growth cabinet (22 °C, 70 % relative humidity, 12 h photoperiod).

### Plant material and growth conditions

Plant lines used in this study are Col-0, *fer-4 (Duan et al., 2010), lrx5x (Herger et al., 2020)* and *ralf7x (Leicher et al., 2026)*.. All plants were grown with two plants per pot in environmentally controlled conditions (21 °C, 55 – 65% relative humidity, 8 h photoperiod). 5-week-old plants were used for *Ecr* infection. Uninfected control samples were transferred to the infection cabinet at the same time as the infected samples, but without powdery mildew infection.

### Sample collection and RNA extraction

For each time point 4 adult leaves from uninfected and infected plants were harvested per genotype. Leaves were directly frozen in liquid nitrogen. The plant material was ground using a TissueLyser. For RNA extraction, TRIzol reagent (Roche, Switzerland) and purification with Direct-zol™ RNA Miniprep Plus kit (Zymo Research, Germany) with on column DNAse I digestion was used. RNA quality was verified by gel electrophoresis on a 2 % agarose gel and RNA quantity was assessed using a NanoDrop ND-1000 Spectrophotometer and a Qubit fluorometer with RNA BR Assay Kit (Thermo Fisher Scientific, USA).

### Library preparation and sequencing

200 ng of RNA samples that passed the quality control were subsequently used for library preparation with the QuantSeq 3‘ mRNAseq V2 Library Prep Kit FWD with Unique Dual Indices (Lexogen, Austria) following the manufacturer’s protocol (user guide 191UG444V0101). Quantification and quality control of the libraries were performed using a Qubit fluorometer with dsDNA HS Assay Kit (Thermo Fisher Scientific, USA) and a Bioanalyzer 2100 with DNA High Sensitivity Kit (Agilent Technologies, USA). The pooled libraries were sequenced on an Illumina NovaSeq 6000 system with single-end reads and 100bp read length (Illumina, USA), aiming for 5-10 M reads per sample.

### Quality Control and Read Trimming

Raw sequencing reads from 80 *Arabidopsis thaliana* RNAseq libraries were assessed for quality using FastQC (v0.12.1) (Andrews, 2010). Adapter sequences and low-quality bases were removed using Trimmomatic (v0.39) (Bolger *et al*., 2014) in single-end mode with the following parameters: ILLUMINACLIP:2:30:10, LEADING:3, TRAILING:3, SLIDINGWINDOW:4:15, MINLEN:40, using a custom adapter file. Trimmed reads were re-evaluated with FastQC.

### Read Alignment

Trimmed reads were aligned to the TAIR10 reference genome (Lamesch *et al*., 2012) using HISAT2 (v2.0.6) (Kim *et al*., 2019). Alignments were performed with default HISAT2 parameters with per-sample read group metadata. Resulting alignments were coordinate-sorted and indexed in csi format using samtools (v0.3.3) (Li *et al*., 2009).

### Read Quantification

Gene-level read counts were generated using feature Counts (subread v2.0.6) (Liao *et al*., 2014) against a modified Araport11 GFF3 annotation file (Cheng *et al*., 2017). A custom script was used to extend the last exon by 3 kb or until the next gene to increase the read count for 3’ mRNA sequencing. Reads were assigned to genes using the ID attribute at the gene feature level with strand-specificity enforced (-s 1). Multi-mapping reads were retained and reads overlapping multiple features were counted for all overlapping features (-M -O).

### Differential Expression Analysis

Prior to differential expression analysis, three samples were identified as outliers based on Euclidean distance from group centroids in PCA space exceeding the 95th percentile of the within-group distribution, confirmed by hierarchical clustering of sample-to-sample distances. These samples were excluded prior to further analysis. Differential expression analysis was performed using DESeq2 (v1.36.0) (Love *et al*., 2014) with a model design of ∼ 0 + Group. Genes with fewer than 10 summed counts across all samples were excluded prior to model fitting. Multiple testing correction was performed using the Benjamini-Hochberg procedure.

### RNAseq analysis

Genes were considered as differentially expressed based on the following criteria: Adjusted p-value ≤ 0.05 and (log2 fold-change) > 1 (upregulated) or (log2 fold-change) < -1 (downregulated). Gene ontology (GO) enrichment was performed using the R package clusterProfiler (v4.14.6) (Yu, 2024) and GO enrichment for Biological Processes (BP) was conducted using the enrichGO function, with gene identifiers specified as TAIR IDs and annotations obtained from the org.At.tair.db database (v 3.20.0) (Carlson, 2024). The background gene universe was defined as all annotated genes available in the database. For gene clustering z-score-transformed transcript abundance (TPM) values were used. Hierarchical clustering was conducted using Euclidean distance and Ward’s minimum variance method (“ward.D2”) as implemented in pheatmap (v1.0.13) (Kolde, 2025) and genes were grouped into five clusters based on the dendrogram structure.

### Conidiophore counting

Conidiophore production was analysed 5 days after infection with *Ecr*. Four leaves from four individual infected plants were harvested per genotype. EtOH:Acetic Acid (6:1) was used to destain the leaves and fungal structures were stained using ink (ink: 25 % acetic acid, 9:1). Conidiophores of single colonies were counted after microscopy using the Zeiss AXIO imager Z1.m microscope with a 20 x magnification.

### Generation of CRISPR-Cas9 mutants

To generate CRISPR-Cas9 mutants of *CYP81F2* the software tool chopchop (https://chopchop.cbu.uib.no/) was used to design 2 target sites (target site 1: AGGGCTTCAATCTCCCACC, target site 2: TATTGTCCGCATGGTCACA). Individual guide RNAs with gene specific target sites were obtained by gene synthesis (Twist Bioscience, USA). The guide RNA fragments were assembled using a GoldenGate-adapted pUC18-based vector. Higher order gRNA stacks were cloned into pICSL4723OD with FastRed-pRPS5::Cas9 (Castel *et al*., 2019). Finally, plant expression constructs were transformed into *Agrobacterium tumefaciens* strain GV3101 for floral dip transformation of Arabidopsis.

## Results

### Powdery mildew infection upregulates the expression of defense genes and downregulates photosynthesis related genes

We used RNAseq to compare transcriptional responses in *fer*-*4*, *lrx5x* and *ralf7x* with Col-0, before and after powdery mildew infection using the fungus *Erysiphe cruciferarum* (*Ecr*). We harvested samples at one and four days post-infection (dpi). Across all samples, a comparable number of genes was detected as expressed (between 18,956 and 21,696, Fig. S1A, Supplementary Data Table 1), and principal component analysis showed a clear separation between Col-0 and the mutant genotypes at both time points post infection (Fig. S1B). However, separation at 1 dpi was mainly driven by differences among genotypes. At 4 dpi, infected samples were clearly separated from untreated controls (Fig. S1C, D). This was also reflected in the higher number of differentially expressed genes (DEGs) that were detected at 4 dpi (Fig. S1 E). At 1 dpi, 1,462 DEGs were detected in Col-0, 258 in *fer*-4, 185 in *lrx5x*, and none in *ralf7x* (Supplementary Data Table 1). However, at 4 dpi, a significantly higher number of DEGs were detected in all genotypes: 7,280 in Col-0, 4,810 in *fer*-4, 4,836 in *ralf7x,* and 5,434 in *lrx5x* (Fig. S1E, Supplementary Data Table 1). This raises the question of whether FER-RALF-LRX pathway mutants generally respond with a weaker transcriptional response to *Ecr* infection.

Due to the much stronger effect observed at 4 dpi and given that the FER-RALF-LRX pathway affects powdery mildew growth at later stages of infection (Leicher *et al*., 2026), we decided to focus on 4 dpi samples for in-depth analysis. At 4 dpi, all genotypes share 3,103 infection-regulated DEGs (Fig. 1A). This accounts for 42.6% of all DEGs found in Col-0. For the mutant genotypes, this group of DEGs accounts for the majority of DEGs (57% for *lrx5x*, 64% *fer-4* and *ralf7x*). This indicates an overlapping effect of *Ecr* infection on the transcriptome of Col-0 and FER-RALF-LRX pathway mutants.

**Figure 1:**
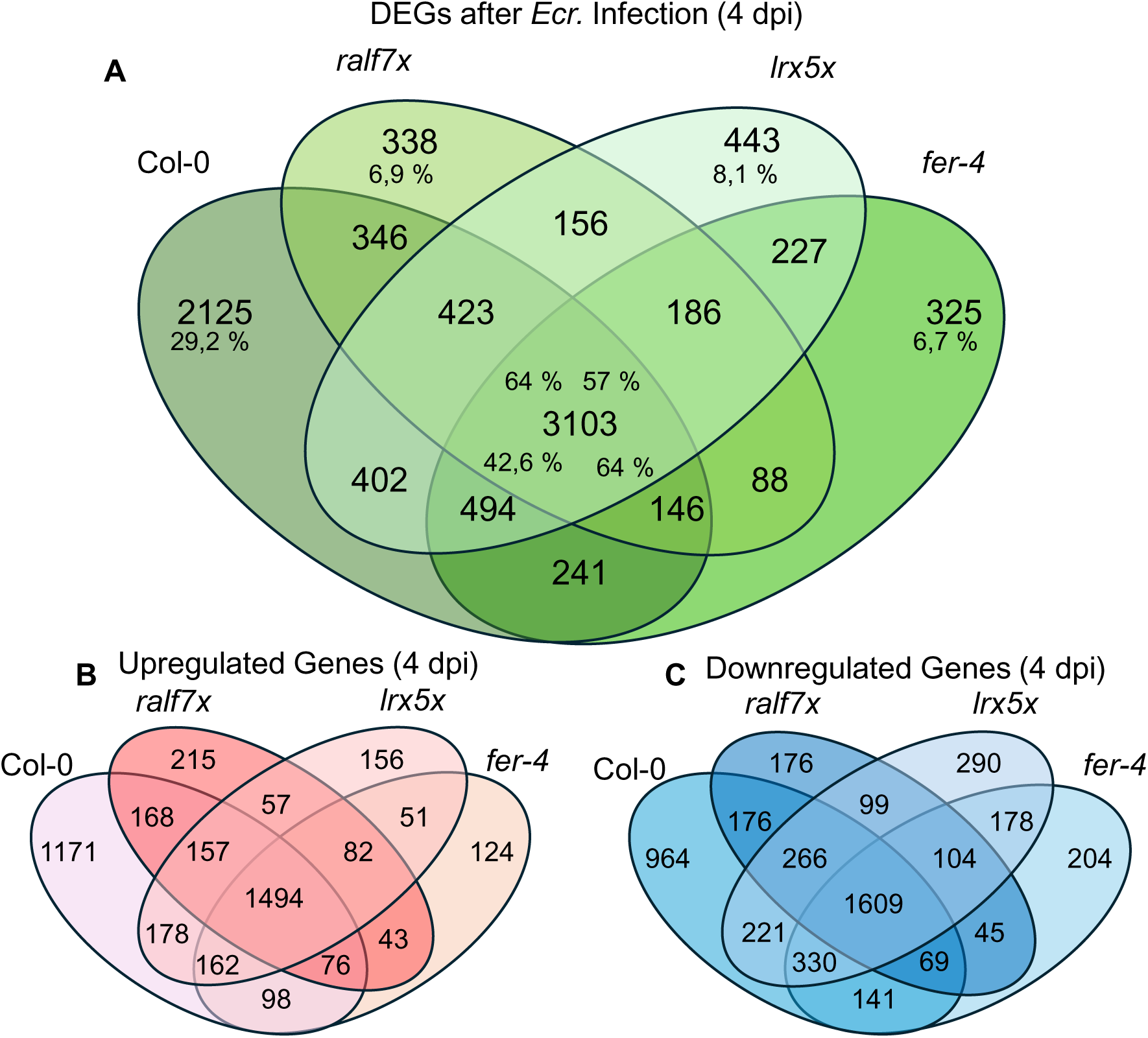
Number of DEGs after *Ecr* Infection in Col-0, *fer-4*, *ralf7x* and *lrx5x*. A) Number of all DEGs after powdery mildew infection (FDR < 0.05, log2(Fold change) > 1or log2(Fold change) < -1). Percentages indicate the proportions of all DEGs of the corresponding genotype located in this field of the Ven-Diagram. B) Number of upregulated genes after powdery mildew infection (FDR < 0.05, log2(Fold change) > 1) C) Number of downregulated genes after powdery mildew infection (FDR < 0.05, log2(Fold change) < -1).

Next, we performed cluster analysis based on the z-score normalized TPM values with all genes that are differentially regulated in at least one genotype at 4 dpi (Fig. 2A, B, Supplementary Data Table 2). Cluster 1 and 2 show increased relative expression in all genotypes after infection compared to the uninfected conditions. Enriched GO-terms in these clusters correlate with response to hypoxia, decreased oxygen levels and immune responses (Fig. 2A, B, Supplementary Data Table 2). Importantly, genes in cluster 2 show similar relative expression values in Col-0 and the mutants before and after infection, indicating largely unaltered general immune responses to *Ecr* infection in FER-RALF-LRX-pathway mutants. The upregulation of defense related genes upon powdery mildew infection, e.g. *PAD4*, *EDS1* or *EDS5* is consistent with previous reports and not strongly affected in our mutants (Fig. 3 A) (Chandran *et al*., 2010). Cluster 3 represents genes with lower relative expression after infection in all genotypes and is enriched in photosynthesis-related genes (Fig. 2A, B, Supplementary Data Table 2). This is most likely linked to changes in resource allocation and a shift from source to sink tissue to exploit host metabolism and support fungal nutrient uptake (Eichmann & Hückelhoven, 2008; Chandran *et al*., 2010). For example, powdery mildew infection induces local upregulation of invertases and sugar transporters and increased cellular sugar content suppresses photosynthesis (Fotopoulos *et al*., 2003; Swarbrick *et al*., 2006). Consistently, we also observed the upregulation of several CW-INVERTASES (CWINV) (*CWINV5*, AT-β-D-FRUCTOFURANOSIDASE 1/*ATBFRUCT1* and *CWINV6*) in all genotypes (Fig. 3B). In addition, SUGAR TRANPORT PROTEIN 4/STP4, a plasma membrane-localized sugar symporter, was previously reported to be upregulated during powdery mildew infection, which we confirm in our dataset (Fotopoulos *et al*., 2003) (Fig. 3B). Importantly, similar to immune responsive genes, the relative expression levels of photosynthesis related genes are comparable between Col-0 and the tested mutants. By contrast, cluster 5 genes show relatively higher normalized expression in all tested mutants compared to Col-0 under control conditions (Fig. 2A). *Ecr* infection leads to cluster 5 downregulation, however, normalized expression values remain relatively higher in the mutants. Most genes of cluster 5 are correlated to wounding, JA signalling and fatty acid responses (Fig. 2B). By contrast, cluster 4 is composed of *Ecr*-induced downregulated genes with relatively lower normalized basal expression in FER-RALF-LRX mutants (Fig. 2A, Supplementary Data Table 2) and is enriched in genes for sulfur-containing glucosinolate metabolism, chloroplast organization, microtubule-based processes and plant cell wall organization/biogenesis (Fig. 2B, Supplementary Data Table 2).

**Figure 2:**
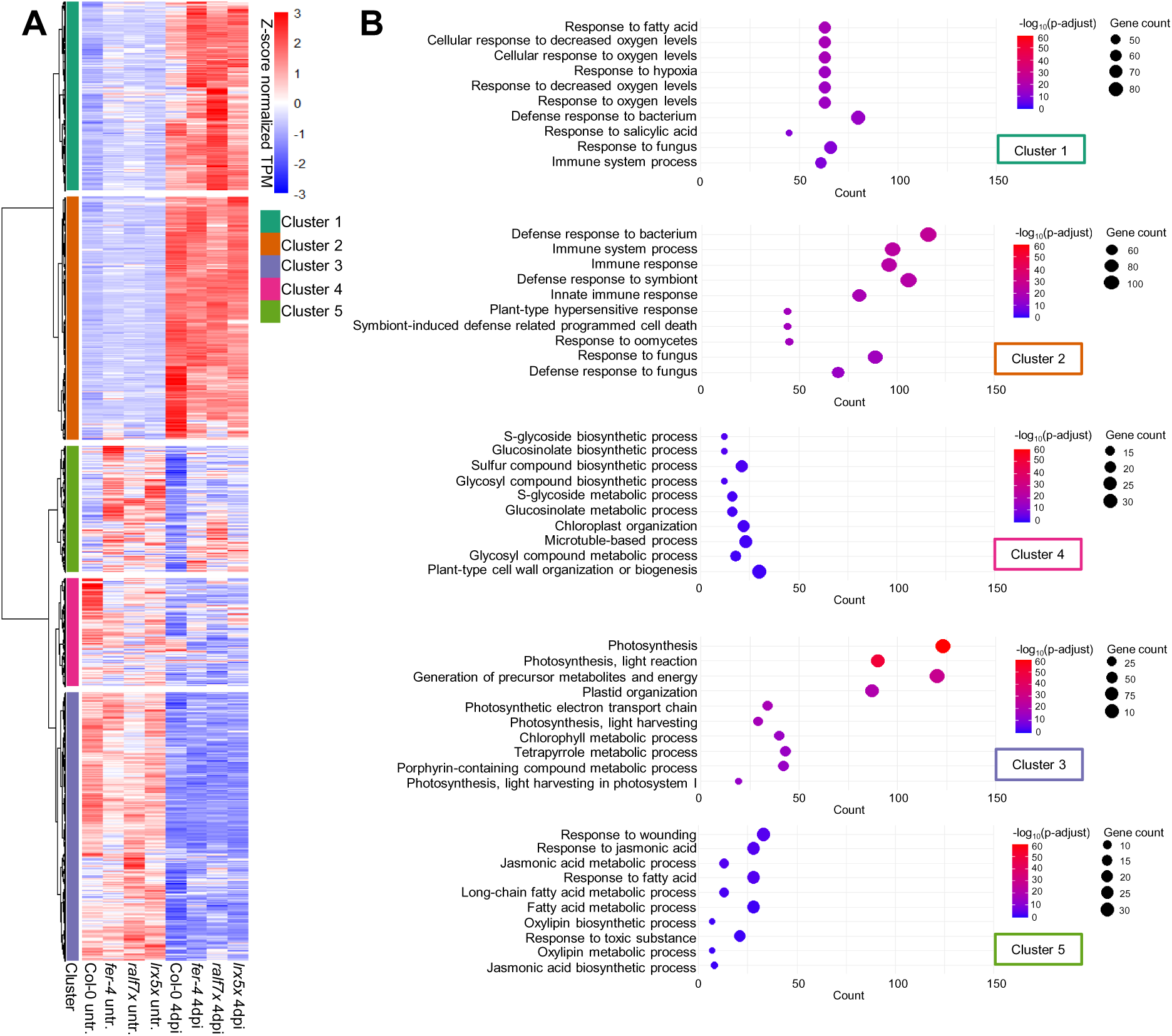
Cluster analysis of all DEGs that are found in at least one genotype at 4 dpi with powdery mildew. A) All DEGs that were found in at least one genotype at 4 dpi post infection were clustered using Euclidean distance and Ward’s minimum variance method in 5 groups according to their z-score normalized TPM-values. B) GO-Term analysis (Biological process) of genes found in the corresponding cluster.

**Figure 3:**
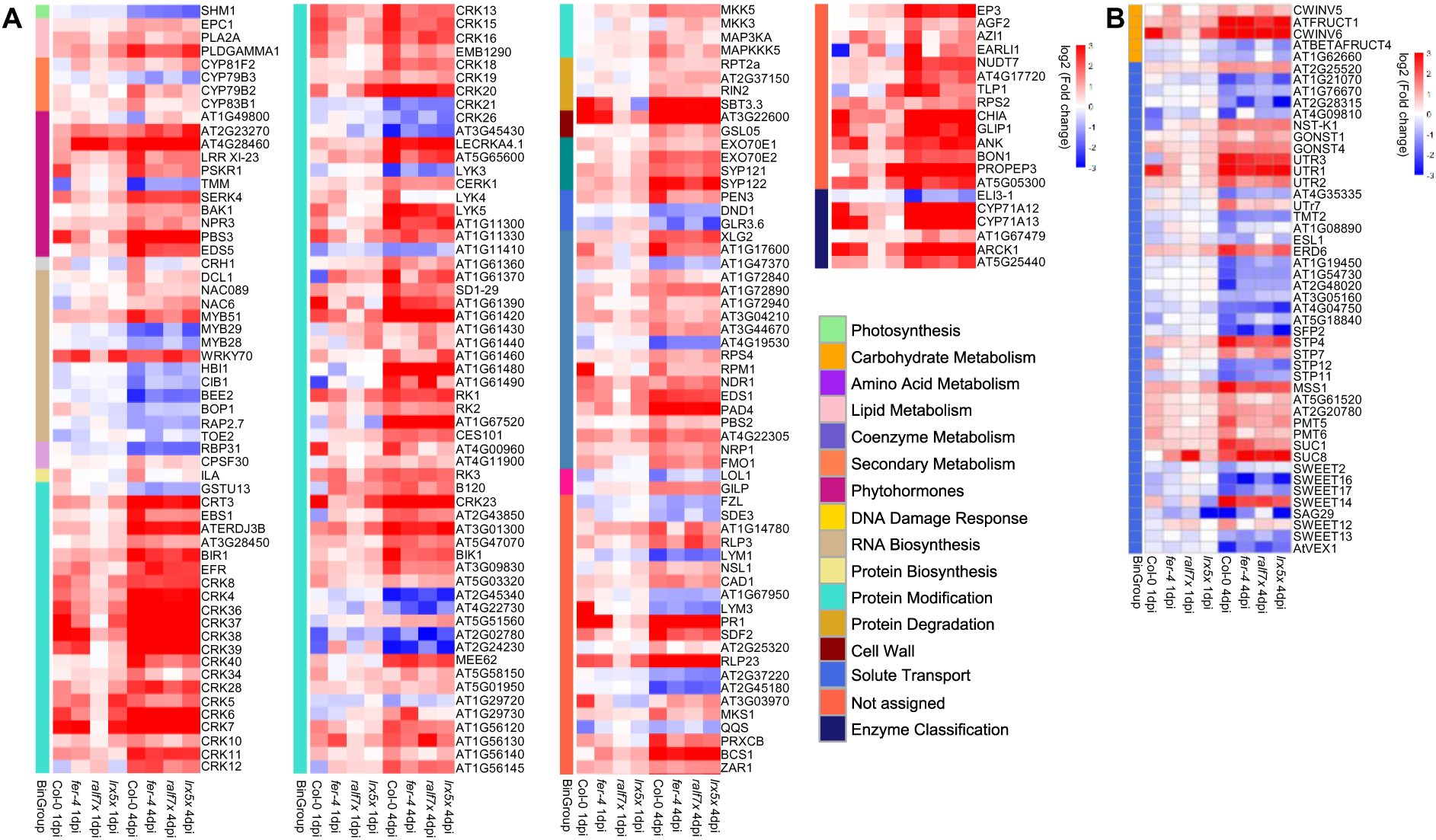
Powdery mildew infection leads to upregulation of defense related genes and differential expression of cwInvertases and transport proteins. A) log2 (Fold change) values (compared to uninfected controls at the same time point) of genes with the GO-term “GO:0006955”: Immune response. All genes that are significantly regulated at 4 dpi with powdery mildew in any of the tested genotypes were considered B) log2(Fold change) of cwInvertases and solute transport proteins at 4 dpi. A) and B) Colour code indicates MapMan Bingroups.

### *Ecr*-induced downregulation of starch metabolism is pronounced in FER/RALF-signalling mutants

Analysis of all *Ecr*-induced DEGs did not reveal clusters that are specifically regulated in Col-0 or the FER-RALF-LRX pathway mutants. To further explore genotype-dependent responses, we performed a more detailed analysis of individual DEGs that are only significant in Col-0, or significant in all mutants but not in Col-0. These may indicate processes that modulate powdery mildew growth on the different host genotypes. No significant GO terms were found within genes specifically downregulated in Col-0 upon infection. DEGs specifically upregulated in Col-0 revealed an enrichment in protein transport and intracellular transport processes (Fig. 4A, Supplementary Data Table 3). Next, we analyzed DEGs that are found in all mutants but not in Col-0, since they could indicate pathways underlying the enhanced resistance of these genotypes. However, GO-term analysis of mutant-specific upregulated genes yielded no significant results. Among mutant-specific downregulated DEGs, genes associated with sugar and starch metabolism were enriched (Fig. 4B, Supplementary Data Table 3).

**Figure 4:**
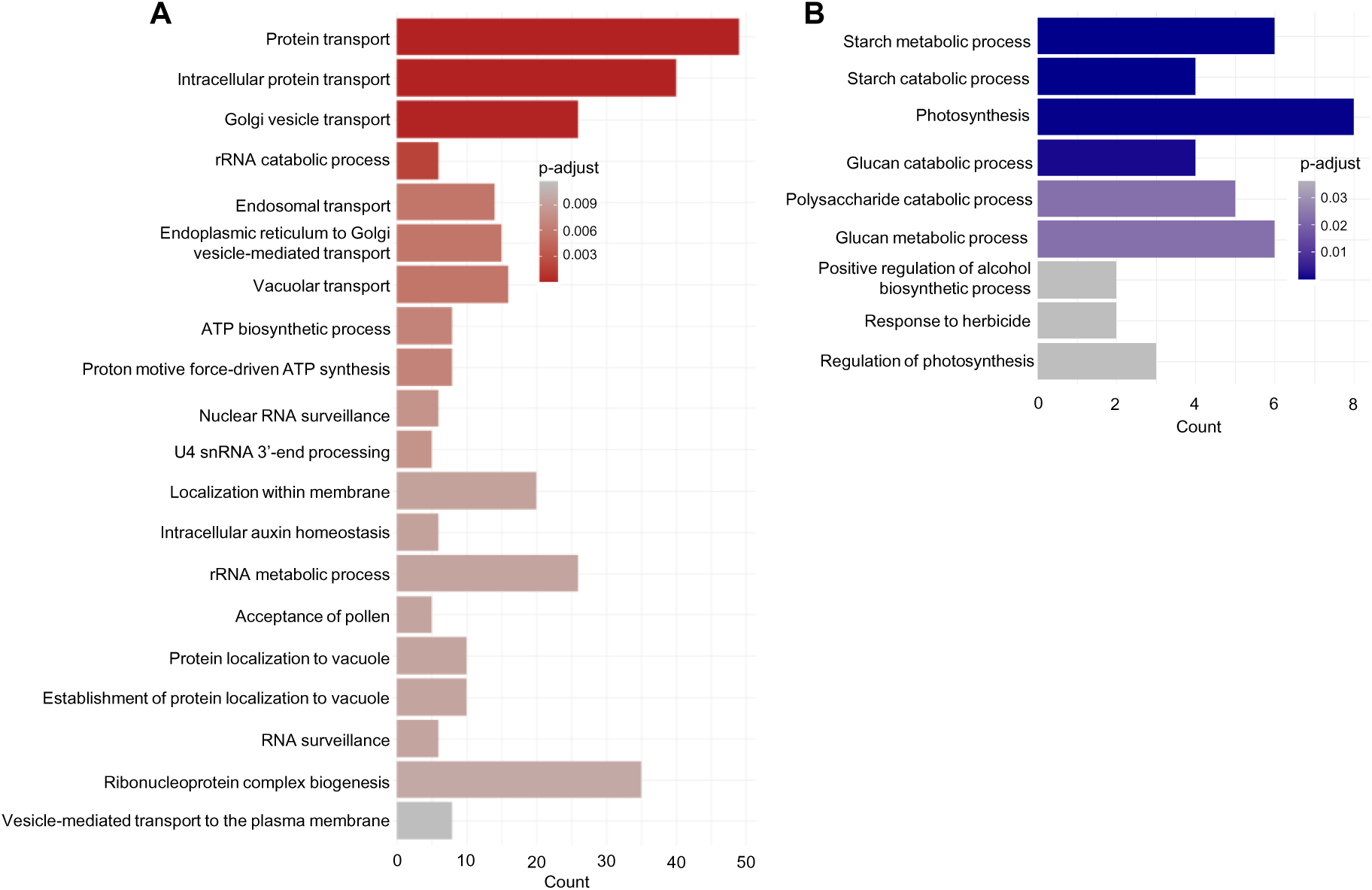
Powdery mildew infection leads to Col-0 specific upregulation of intracellular transport processes and pronounced effects on starch metabolism in FER-RALF-LRX pathway mutants. A) GO-Term analysis (Biological Process) of all DEGs that are specifically upregulated only in Col-0 at 4 dpi. B) GO-Term analysis (Biological Process) of all DEGs that are specifically downregulated in all mutants but not found in Col-0.

Collectively, the general response to powdery mildew infection does not appear to strongly differ between Col-0 and the FER-RALF-LRX-pathway mutants. All genotypes show strong upregulation of defense related genes and downregulation of photosynthesis, indicating a shift from source to sink tissue. However, specific analysis of DEGs that are exclusively regulated in Col-0 or in the mutants revealed an upregulation of intracellular transport processes in Col-0 and a stronger downregulation of starch and sugar metabolism in the mutant genotypes. Further, cluster analysis identified genes that do not show differences in the regulation of gene expression during infection but display higher or lower expression levels in the untreated mutant plants compared to Col-0 (Fig. 2A).

### FER-RALF-LRX affects expression of JA, tryptophan and indol glucosinolate-related genes

Previous work showed that the *fer-4* mutant displays constitutive transcriptional differences compared to the wild type. FER phosphorylates and destabilizes the JA-responsive transcription factor MYC2, explaining constitutive upregulation of JA-associated genes in *fer-4* (Guo *et al*., 2018). Furthermore, genes associated with glucosinolate metabolism and ER body formation show higher steady-state levels in *fer-*4 (Wang *et al*., 2022). Since significant differences in *Ecr*-induced gene expression likely does not explain the enhanced resistance of FER-RALF-LRX pathway mutants and cluster 4 and 5 indicated steady-state differences in relative gene expression levels of several genes (Fig. 2A), we next compared the transcriptional profile of untreated FER-RALF-LRX pathway mutants with Col-0.

PCA component analysis of untreated samples clearly separated the mutant genotypes from Col-0 with a PC1 of 67 % (Fig. 5A). When compared to the wildtype Col-0, we identified 1,205 DEGs in *fer-*4, 1,242 in *ralf7x* and 652 DEGs in *lrx5x* (Fig. 5B, Supplementary Data Table 4). Interestingly, while 30 % and 37 % of DEGs were specific for either *fer-*4 or *ralf7x,* respectively, only 9 % of DEGs were specific for *lrx5x* (Fig. 5C). Assuming a common mechanism for increased resistance in all mutants, we focused our analysis on DEGs shared among all mutants compared to Col-0. 461 DEGs matched this category, with 396 displaying higher and 65 lower transcript abundances compared to Col-0, respectively (Fig. 5C, D, E). We next performed cluster analysis with the z-score normalized TPM values of all genes that are significantly different in all mutant genotypes compared to Col-0 (Fig. 6A, Supplementary Data Table 5). Again, clusters showing relatively higher normalized basal expression levels compared to Col-0 (cluster 3 and 4) were enriched in genes associated with wounding response, fatty acid response and JA signalling. In addition, indole-containing compound metabolic processes and amine biosynthetic processes are enriched in these clusters (Fig. 6A, B, Supplementary Data Table 5). Cluster 2 also contains genes with relatively higher normalized expression in the mutants compared to Col-0, however, the differences in normalized expression levels are less pronounced. Cluster 2 was enriched in tryptophan biosynthetic processes and aromatic amino acid metabolic processes (Fig. 6A, B, Supplementary Data Table 5).

**Figure 5:**
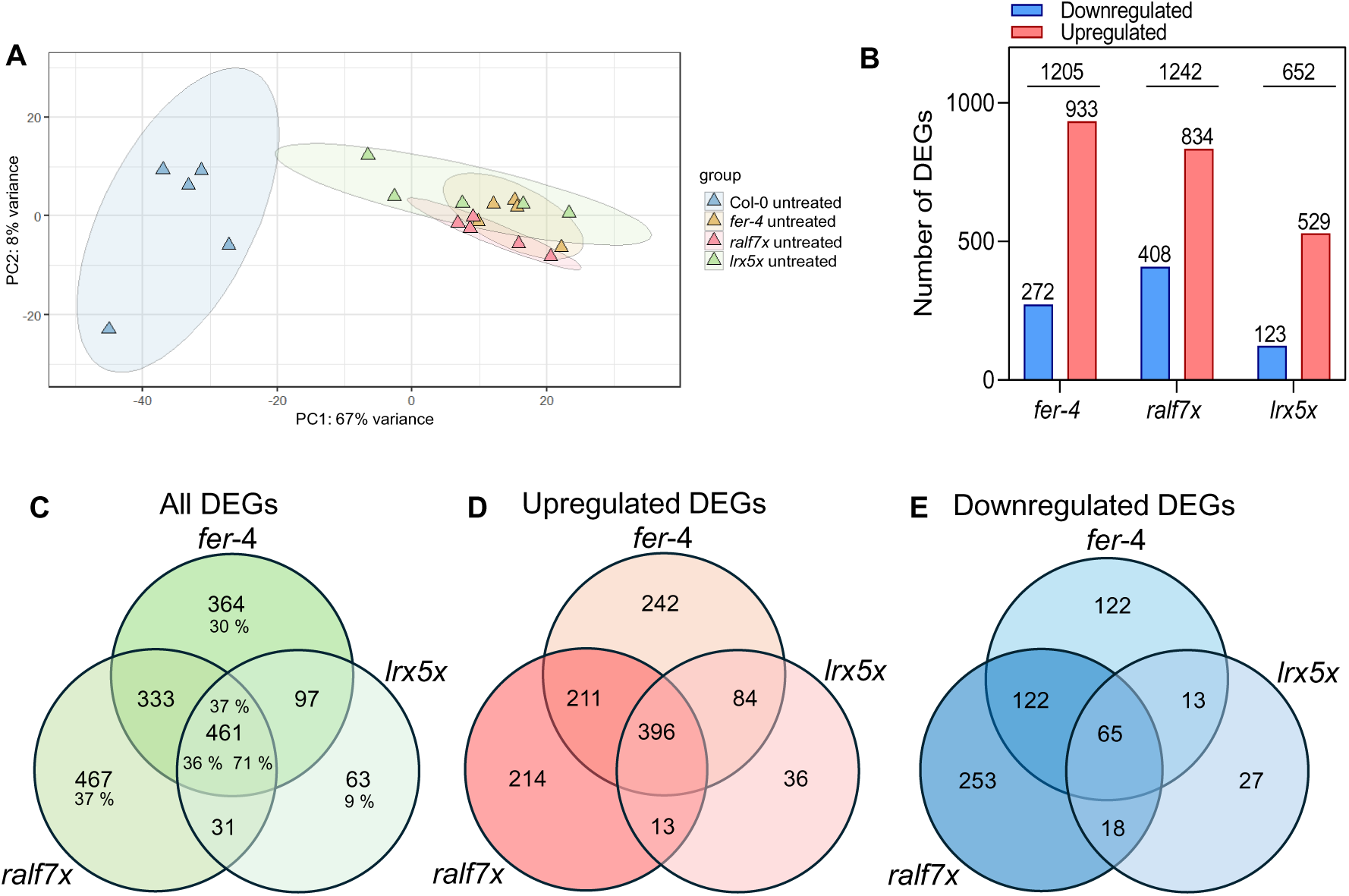
RALF/FER/LRX-signalling mutants show steady-state differences in gene expression at an untreated stage compared to Col-0. A) PCA-Analysis based on the variance stabilized counts from DESeq2. B) Number of DEGs in untreated fer-4, ralf7x and lrx5x compared to Col-0. C) Number of all DEGs found in the untreated plants compared to Col-0 (FDR < 0.05, log2(Fold change) > 1or log2(Fold change) < -1). Percentages indicate the proportions of all DEGs of the corresponding genotype located in this field of the Ven-Diagram. D) Number of upregulated genes found in the untreated plants compared to Col-0 (FDR < 0.05, log2(Fold change) > 1) C) Number of downregulated genes found in the untreated plants compared to Col-0 (FDR < 0.05, log2(Fold change) < -1).

**Figure 6:**
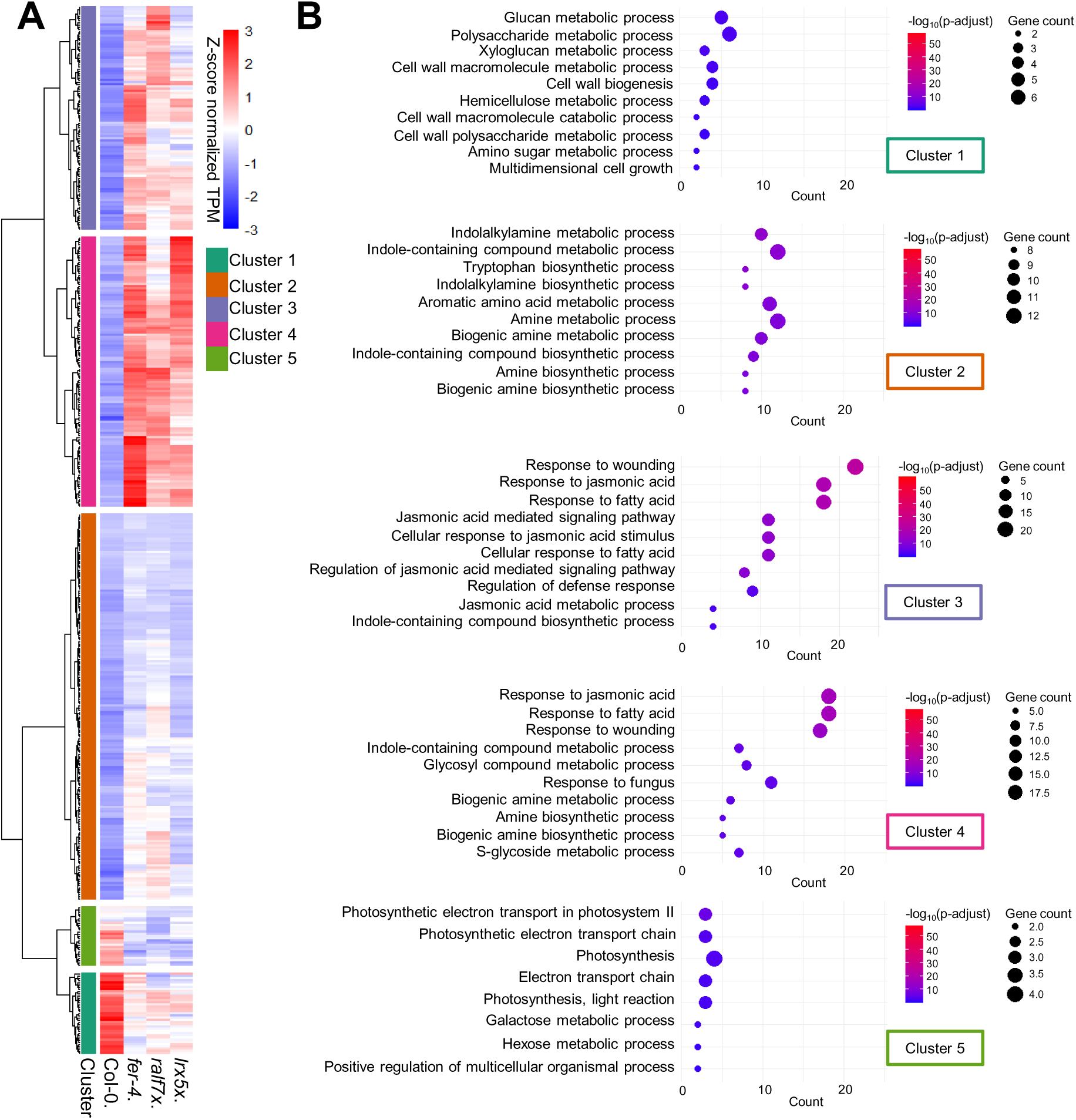
Cluster analysis of all DEGs that are found in at least one genotype in comparison to Col-0 (untreated) A) All DEGs that were found in at least one genotype at an untreated stage compared to Col-0 were clustered using Euclidean distance and Ward’s minimum variance method in 5 groups according to their z-score normalized TPM-values. B) GO-Term analysis (Biological process) of genes found in the corresponding cluster.

We next filtered individual genes correlated with MapMan BINcode 11.7: Phytohormones/JA (Schwacke *et al*., 2019). Enzymes involved in almost every step of JA biosynthesis were upregulated in all three mutants (Fig. 7A, B, Fig. S3A). Furthermore, 239 of the 396 genes upregulated in all three mutants (60.3%) are listed as targets of MYC2 and/or MYC3 (Zander *et al*., 2020) (Supplementary data table 6). JA treatment can induce tryptophan and indole glucosinolate (IGL) biosynthesis (Cao *et al*., 2016). Consistently, we observe a strong upregulation of genes associated with tryptophan biosynthesis in FER-RALF-LRX pathway mutants (Fig. 7C, D, Fig. S3B). Tryptophan serves as a precursor for auxin, camalexin, and IGL biosynthesis. In roots RALF1 treatment increases auxin biosynthesis in a FER- dependent manner through upregulation of *YUC* expression, enzymes involved in auxin biosynthesis (Li *et al*., 2022). However, no enzymes of the auxin biosynthesis pathway were found as significant DEGs in any of the mutants, indicating that knockout of the FER-RALF-LRX pathway does not affect auxin biosynthesis on a transcriptional level in leaves.

**Figure 7:**
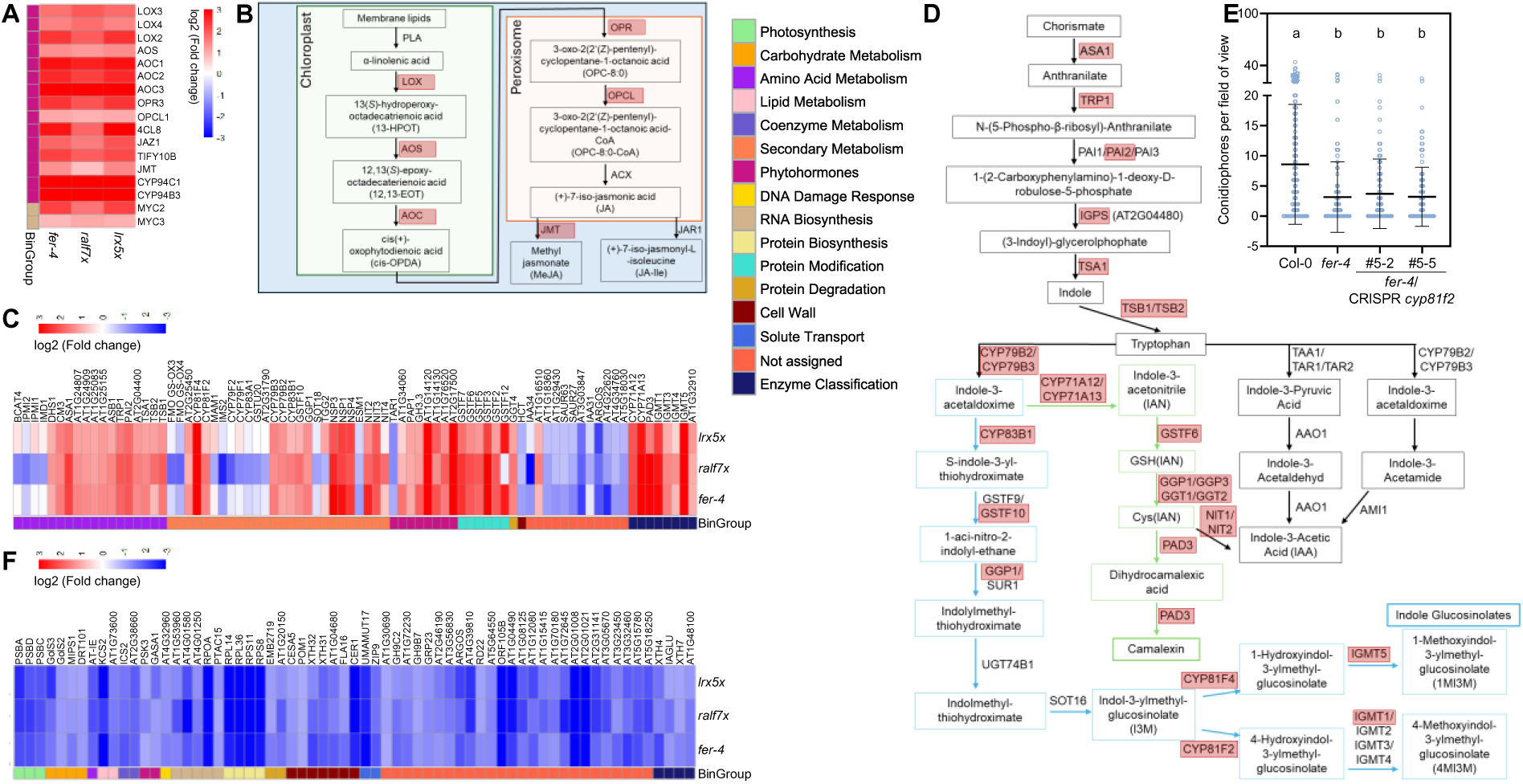
FER-RALF-LRX pathway mutants show steady-state upregulation of genes involved in JA signalling and tryptophan derived secondary metabolism. A) log2(Fold change) values of genes involved in JA biosynthesis and signalling. B) Pathway of JA biosynthesis. Genes for enzymes marked in red were found to be upregulated in FER-RALF-LRX pathway mutants. C) log2(Folg change) values of DEGs correlated to the shikimate pathway, glucosinolate, camalexin and auxin metabolism. D) Pathway of tryptophan biosynthesis and biosynthesis of tryptophan derived secondary metabolites. Genes for enzymes marked in red were found to be upregulated in FER-RALF-LRX pathway mutants. E) Conidiophores per field of view (5 dpi) of fungal colonies grown upon *Ecr* infection of the indicated genotypes. Mean ± SD, n=108-209 pooled from three independent experiments (Dunn’s multiple comparisons test, a-b p≤0.0001). F) log2(Fold change) of DEGs that are downregulated in all mutants genotypes compared to Col-0 at an untreated stage.

By contrast, gene expression of enzymes regulating camalexin and IGL biosynthesis were upregulated in all mutants, similar to previous reports in *fer-4 (Wang et al., 2022)* (Fig. 7C, D). Three indole glucosinolates (indol-3-ylmethyl; I3M), 4-methoxy-indol-3-ylmethyl; 4MI3M), and 1-methoxy-indol-3-ylmethyl; 1MI3M) show increased accumulation in *fer*-4 (Wang *et al*., 2022). CYP81F2 catalyses conversion of I3M to 4MI3M, a process that has been linked to non-host resistance against non-adapted powdery mildew (Fig. 7D) (Hunziker *et al*., 2020). To test whether increased production of I3M/4MI3M explains the resistance of FER-RALF-LRX pathway mutants to *Ecr*, we generated two allelic CRISPR *cyp81f2* mutants in *fer-4* (Fig. S3C). However, CRISPR *cyp81f2* did not rescue the reduced *Ecr* conidiation phenotype of *fer-4* (Fig. 7E). Yet we cannot rule out effects of other accumulating IGLs in FER-RALF-LRX pathway mutants on *Ecr* resistance. For example, CYP81F4, involved in 1MI3M production is similarly upregulated in our mutants and represents an CYP81F2-independent branch of IGL biosynthesis (Fig. 7C, D) (Kai *et al*., 2011).

### Cell wall related genes are downregulated in FER-RALF-LRX signalling mutants

Cluster 1 and cluster 5 of differential steady-state expressed genes between Col-0 and FER-RALF-LRX pathway mutants contains genes that are relatively lower expressed in the mutant genotypes (Fig. 6A, B). Cluster 5 is strongly enriched in GO-terms related to photosynthesis, while genes in cluster 1 are mainly associated with cell wall metabolic processes (Fig. 6B). Among the 65 genes that are significantly downregulated in all mutants compared to Col-0 we identified *CELLULOSE SYNTHASE 5* (*CESA5)* and *POM-POM1* (*POM1*) (Fig. 7F, Fig. S3 D), enzymes that are required for cellulose biosynthesis (Endler & Persson, 2011; Sánchez-Rodríguez *et al*., 2012). This is consistent with reduced cellulose content and altered cell wall monosaccharide composition in *fer-*4 (Yeats *et al*., 2016; Biermann *et al*., 2025). Further, several xyloglucan endotransglucosylase/hydrolase (XTH) proteins are downregulated in all mutants (Fig. 7F, Fig. S3 D). XTHs are cell wall modifying enzymes that can cut, re-join or hydrolyse the hemicellulose component xyloglucan and by this modulate cell wall plasticity and cell expansion (Ishida & Yokoyama, 2022). Changes in xylan, xyloglucan content and xyloglucan acetylation have been linked to immunity against several plant pathogens, including powdery mildew (Delgado-Cerezo *et al*., 2012; Chowdhury *et al*., 2017). However, whether XTH activity directly affects powdery mildew infection success remains unknown.

### Analysis of genes with differential expression in Col-0 upon *Ecr* infection and altered transcript accumulation in FER-RALF-LRX pathway mutants

Comparison between the transcriptome of Col-0 to *fer-*4, *ralf7x* and *lrx5x* revealed severe changes in JA, cell wall, tryptophan and IGL-related gene expression. In a next step, we searched for genes that are mis-regulated in all mutant genotypes and, in addition, differentially expressed in Col-0 after powdery mildew infection. We found 284 genes that fit these criteria (Fig. 8A). GO-term analysis showed that genes involved in JA signalling, response to fatty acid and wounding, as well as indole alkylamine metabolic and tryptophan biosynthetic processes were enriched, confirming our previous analysis (Fig. S4A). Upon cluster analysis of these genes, we identified one cluster (cluster 1) with genes displaying relatively stronger expression in Col-0 under control conditions. The majority of those genes were downregulated in all genotypes upon *Ecr* infection (Fig. 8 B, C, Fig. S4B). This cluster contained genes involved in cell wall processes, including CESA5 and POM1 and two XTH genes (XTH32, XTH31), as well as the histidine biosynthesis enzyme AT-IE/HISTIDINE BIOSYNTHESIS 2 (AT-IE/HISN2). Genes in cluster 2 and 3 showed relatively higher basal expression in our mutants and were upregulated by *Ecr* infection in all genotypes (Fig. 8C, D, Fig. S4B). By contrast, genes in cluster 4 and 5 showed normalized steady-state expression levels that were higher in our mutants but were normalized expression values were downregulated after *Ecr* infection in all genotypes (Fig. 8C, D, Fig. S4B). Consistent with our previous analysis, we identified JA-related genes (cluster 2), tryptophan biosynthesis (cluster 2) and IGL biosynthesis enzymes (cluster 3) among those genes. However, the direction of regulation appears similar for the vast majority of genes in all 5 clusters, raising the question whether dose-dependent metabolic shifts related to secondary metabolism, hormone homeostasis and cell wall remodelling may explain the resistance phenotype of FER-RALF-LRX pathway mutants. Collectively, these genes may represent physiologically relevant targets of the FER-RALF-LRX pathway to support fungal accommodation.

**Figure 8:**
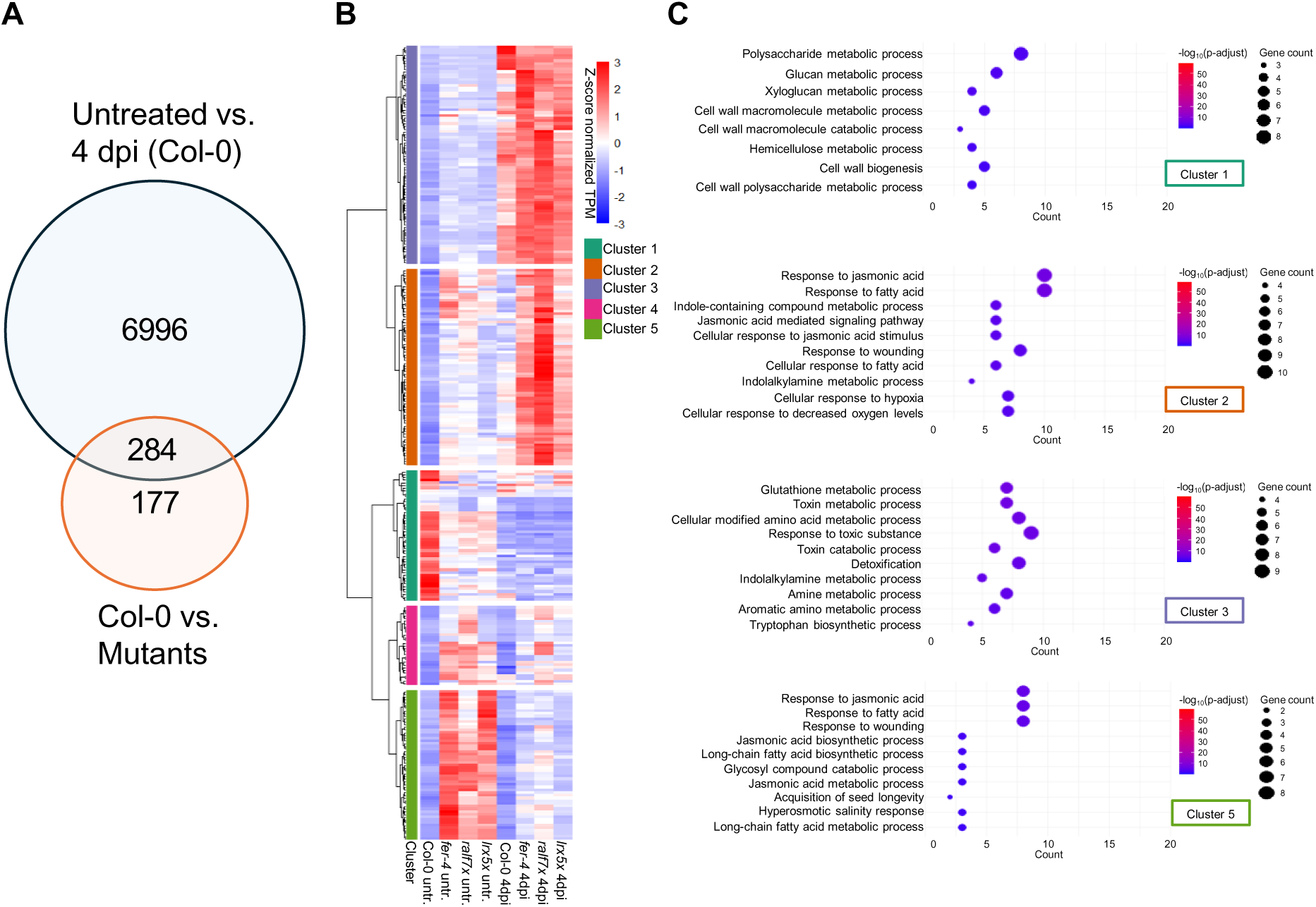
Cluster analysis of all DEGs that are found in Col-0 at 4 dpi and in all mutants compared to Col-0 at an untreated stage. A) Ven-Diagram of all DEGs that are found in Col-0 at 4 dpi and in all mutants compared to Col-0 at an untreated stage. B) All DEGs that are found in Col-0 at 4 dpi and in all mutants compared to Col-0 at an untreated stage (284 genes from A) were clustered using Euclidean distance and Ward’s minimum variance method in 5 groups according to their z-score normalized TPM-. C) GO-Term analysis (Biological Process) of DEGs within in the corresponding cluster.

## Discussion

This study investigates the transcriptional response of Col-0 and FER-RALF-LRX-signalling mutants to powdery mildew infection. We analysed gene expression data at 1 and 4 dpi and compared the transcriptomes of untreated plants to understand the effects of powdery mildew infection and to shed light on infection-independent differences in mature rosette leaves. Our results suggest that FER-RALF-LRX signalling does not significantly impact the primary responses to powdery mildew infection, such as enhanced defense gene expression or the transition from source to sink tissue. Instead, FER-RALF-LRX-dependent effects on metabolism and cell wall-related genes may be associated with reduced conidiation in these mutants.

A higher number of DEGs were found in Col-0 after powdery mildew infection compared to the tested mutants. While this may be related to their reduced infection level (Leicher *et al*., 2026), the general responses were comparable between all tested genotypes. As expected, infection with powdery mildew led to the upregulation of defense-related genes. However, the changes in expression of biotic stress genes was not significantly different between FER-RALF-LRX pathway mutants and Col-0 (Fig. 3A). This suggests that the mutants’ resistance phenotype is not based on generally elevated immune responses and is consistent with *fer*-4 and *ralf7x* being more susceptible to other pathogens, such as *Pseudomonas syringae* DC3000 pv. *tomato* or *Hyaloperonospora arabidopsidis* (Stegmann *et al*., 2017; Leicher *et al*., 2026).

During infection, photosynthesis-related genes were downregulated in all genotypes, which is in accordance with previous reports investigating various powdery mildew species on different hosts (Swarbrick *et al*., 2006; Li *et al*., 2020). Powdery mildew fungi fully rely on nutrient uptake from their hosts (Hückelhoven & Panstruga, 2011) and thus likely actively induce metabolic shifts in infected cells (Wright *et al*., 1995). This is accompanied by increased cell wall invertase activity and the upregulation of hexose transporters (Wright *et al*., 1995; Fotopoulos *et al*., 2003; Sutton *et al*., 2007; Chandran *et al*., 2010). Accordingly, we also observe the upregulation of cell wall invertases (ATBFRUCT1, CWINV5, CWINV6) and sugar transporters, such as STP4 and SUC1 (Fig. 3B). Cell wall invertases typically operate in phloem unloading by cleaving apoplastic sucrose into glucose and fructose, which are then transported into the cell by sugar transporters (Roitsch & González, 2004; Proels & Hückelhoven, 2014). Sugar cleavage by CW-invertases may be necessary since it has been reported that wheat powdery mildew prefers glucose over sucrose as a sugar source (Sutton *et al*., 2007). In addition, enhanced levels of soluble sugars in the apoplast may inhibit the export of hexoses from the infected cell. Increased sugar levels inhibit photosynthesis, resulting in decreased gene expression of photosynthesis-related genes, consistent with our analysis (Berger *et al*., 2007).

A notable difference between Col-0 and FER/-RALF-LRX signalling mutants was a stronger downregulation of starch metabolism-related genes upon *Ecr* infection (Fig. 4B, Fig. S2D). FER has previously been linked to starch metabolism, with *fer* mutants accumulating higher starch levels upon sucrose treatment (Yeats *et al*., 2016). This phenotype was attributed to increased apoplastic acidification, which induces enhanced sucrose uptake (Yeats *et al*., 2016). Furthermore, FER interacts with the key cytosolic glycolytic enzyme glyceraldehyde-3-phosphate dehydrogenase (GAPDH), and FER deficiency results in reduced GAPDH activity and increased starch accumulation (Yang *et al*., 2015). During conidiation of *Blumeria graminis f. sp. tritici* (wheat powdery mildew), fungal genes involved in converting sucrose to glucose, glycolysis and the tricarboxylic acid cycle, as well as other pathways related to energy metabolism, are activated (Zeng *et al*., 2017). This supports a high-energy demand required for this process. Wheat infection with powdery mildew leads to reduced starch accumulation in grains by repressing enzymes related to starch synthesis (Gao *et al*., 2014). The fungus may actively repress starch biosynthesis to enhance cellular soluble sugar content. Although starch synthesis genes were also downregulated in Col-0, this effect was more pronounced in the FER-RALF-LRX pathway mutants (Fig. S2B). This raises the possibility that enhanced starch biosynthesis in *fer-4*, *lrx5x* and *ralf7x* requires stronger repression by the fungus to shift the balance from starch to sucrose. However, starch-free mutants are less susceptible to *Ecr (Engelsdorf et al., 2013)*. Hence, mutation of starch synthesis genes in FER-RALF-LRX pathway mutants could help to clarify how this pathway may shape starch metabolism to influence powdery mildew susceptibility.

FER-RALF-LRX signalling regulates multiple aspects of plant physiology. In line with that, strong transcriptomic differences were already evident in untreated samples. Analysis of downregulated DEGs in all mutants revealed an enrichment of genes involved in cell wall metabolic processes (Cluster 1 in Figure 6A, B). The cell wall of *fer-*4 has reduced cellulose levels (Yeats *et al*., 2016; Biermann *et al*., 2025). In roots FER interacts with CELLULOSE SYNTHASE (CESA)-INTERACTIVE PROTEIN 1 (CSI1), a protein required for cellulose biosynthesis (Yu *et al*., 2024). Cellulose is synthesised at the plasma membrane by cellulose synthase (CESA) complexes (CSCs). Interestingly, *cesa3* displays reduced cellulose levels and, upon powdery mildew infection, shows reduced fungal conidiation and elevated JA signalling, reminiscent of *fer-4* (Ellis & Turner, 2001; Leicher *et al*., 2026). Cellulose microfibrils are embedded in a network of hemicellulose and pectin (Cosgrove, 2024). The most abundant hemicellulose is xyloglucan, and genes related to hemicellulose metabolism, in particular xyloglucan metabolism, are downregulated in FER-RALF-LRX pathway mutants. XTH genes mediate xyloglucan remodelling, which affects cell wall stiffness (Rose *et al*., 2002). *Marchantia polymorpha fer* mutants also show reduced transcript accumulation of genes linked to xyloglucosyl transferase activity, many of which were also differentially regulated in Col-0 upon *Ecr* infection. Immunolabelling experiments in *Marchantia polymorpha* suggested changes in either the xyloglucan content or its accessibility in the cell wall of *fer* mutants (Muller *et al*., 2025). However, the effect of changes in xyloglucan levels or modification on powdery mildew colonisation is currently unknown. Changes in cell wall composition may interfere with the activity of cell wall invertase (cw-INV) and alter sugar availability in the apoplast, potentially leading to nutritional defects in the fungus. Therefore, combined with the observed differences in starch metabolism in the mutants, further analysis of cell wall and intracellular sugar content, with and without fungal infection, is required to fully understand potential FER-RALF-LRX-dependent alterations in sugar metabolism.

Cell wall perturbations are tightly linked to JA-signalling (Mielke & Gasperini, 2019). Cell wall damage alone can induce JA biosynthesis, and FER negatively regulates JA-signalling through interaction with and destabilization of the transcription factor MYC2 (Guo *et al*., 2018). Among DEGs shared between all mutants, 55% were annotated as MYC2- and/or MYC3-dependent (Zander *et al*., 2020) (Supplementary data table 6). This indicates a major contribution of JA-signalling to the mutants’ transcriptomes. While enhanced JA-signalling in *fer* mutants has been attributed to loss of FER-MYC2 interaction (Guo *et al*., 2018), our data raises the question whether persistent cell wall integrity signalling also contributes to constitutive JA activation in FER-RALF-LRX pathway mutants (Bacete *et al*., 2022). However, our previous analysis showed that mutating *MYC2* in *fer-4* does not affect *Ecr* resistance, suggesting that constitutive MYC2-dependent JA signalling does not explain this phenotype (Leicher *et al*., 2026). Enhanced JA-signalling promotes tryptophan biosynthesis (Johnson *et al*., 2023). Consistent with this, we observe increased expression of those genes in all genotypes, many of which were also differentially expressed in WT upon *Ecr* infection. Similarly, genes associated with IGL and camalexin biosynthesis are upregulated in FER-RALF-LRX mutants. Upregulation of IGL biosynthesis has previously been reported in *fer-*4, accompanied by increased levels of I3M, 4MI3M and 1MI3M (Wang *et al*., 2022). IGL biosynthesis has notably been associated with abaxial leaf immunity against powdery mildew infection (Wu *et al*., 2023). Furthermore, the IG 4MI3M is required for non-host resistance. Infection with non-adapted powdery mildew leads to the accumulation of 4MI3M, and mutation of *CYP81F2* (which is also upregulated in all tested mutants) results in reduced 4MI3M levels and impaired penetration resistance against non-adapted powdery mildew fungi (Bednarek *et al*., 2009; Hunziker *et al*., 2020). However, mutation of *CYP81F2* in *fer-*4 mutant did not restore powdery mildew susceptibility, indicating that enhanced resistance in the mutant is independent of 4MI3M (Fig. 7E). Nevertheless, it cannot be excluded that the accumulation of other antifungal precursors or enhanced 1MI3M production masks the effect of reduced 4MI3M levels. Mutational analysis of additional IGL biosynthesis enzymes could clarify their contribution to the FER-RALF-LRX mutant resistance phenotype.

Camalexin is another tryptophan-derived antimicrobial compound with upregulated expression of biosynthesis genes in FER-RALF-LRX pathway mutants. Enhanced accumulation of camalexin leads to increased resistance against the powdery mildew fungus *G. cichoracearum (Liu et al., 2016)*. However, no enhanced camalexin levels were detected in *fer-4* or *ralf7x* before or after powdery mildew infection (Leicher *et al*., 2026). In addition, camalexin increases resistance to bacterial and oomycete infections, but *fer-4* is more susceptible to both pathogens (Stegmann *et al*., 2017; Lopa *et al*., 2025; Leicher *et al*., 2026). Therefore, the *Ecr* resistance phenotype in FER-RALF-LRX pathway mutants is most likely independent of camalexin.

FER activates the TARGET OF RAPAMYCIN (TOR), a central growth regulator that coordinates growth with nutrient and energy status (Song *et al*., 2022). Further, FER regulates salt stress tolerance through phosphorylation and stabilization of SERINE HYDROXYMETHYLTRANFERASE 1, which modulates photorespiratory flux (Jiang *et al*., 2024). Both studies reported altered amino acid profiles in *fer-4*. The enhanced expression of tryptophan biosynthesis genes prompted us to investigate whether other genes involved in amino acid metabolism are differentially expressed in the FER-RALF-LRX pathway mutants. However, tryptophan biosynthesis was the only amino acid pathway with several differentially regulated genes. Interestingly, AT-IE/HISN2, is the only gene that is attributed to amino acid metabolism with lower basal expression in all mutants. *Ecr* infection reduced *HISN2* expression across all genotypes, with lower infection-induced abundance in FER-RALF-LRX pathway mutants. HISN2 is a bifunctional enzyme and catalyses the second and third steps of histidine biosynthesis (Ingle, 2011). Overexpression of an *LRX1* variant lacking the extensin domain (LRX1ΔE14) causes a dominant-negative effect on root hair formation, which can be suppressed by mutation of *HISN2* (Guérin *et al*., 2024). So far, analysis of the amino acid profile in *fer-4* revealed higher levels of Histidine compared to the wildtype, which does not correlate with our reported downregulation of *HISN2 (Song et al., 2022; Jiang et al., 2024)*. Yet, these measurements were performed in seedlings while our gene expression data was obtained from rosette leaves of adult plants, raising the question of age-dependent differences for FER-RALF-LRX-regulated amino acid biosynthesis gene expression.

Collectively, our data supports a picture, in which FER-RALF-signalling mutants retain a largely intact canonical defense response to powdery mildew but exhibit altered cell wall composition, sugar metabolism, and JA-dependent secondary metabolism. Resistance of *fer-4* and *ralf7x* is characterized by normal penetration success but strongly impaired conidiophore production, suggesting that fungal reproduction rather than initial infection is compromised (Leicher *et al*., 2026). We propose that changes in cell wall polysaccharides and carbon allocation restrict nutrient availability to the fungus, leading to starvation during the energy-intensive conidiation phase. Future studies should focus on detailed analyses of cell wall composition and monosaccharide profiles in FER-RALF-LRX-signalling mutants, both under control conditions and during infection. Metabolomic profiling of sugars and secondary metabolites, combined with feeding experiments, may help to elucidate nutrient fluxes from host to pathogen to understand metabolic changes caused by powdery mildew infection in a FER-RALF-LRX-dependent manner. Such approaches will be essential to better understand the mechanistic basis underlying FER-LRX-RALF-dependent powdery mildew susceptibility.

## Acknowledgements

This work was funded by the Deutsche Forschungsgemeinschaft (DFG) grants 465669230 (STE2448/4-1 to MS), 491090170 (TRR356/1 TP B09 (to MS), Z03 (to NK) and TP A03 (to RH)). The work was further supported by core funding from the Technical University of Munich (MS, RH, NK) and Ulm University (MS). We thank Stefanie Ranf for providing GoldenGate vectors for molecular cloning and Mark Youles (TSL Norwich) and Laurence Tomlinson (TSL Norwich) for providing the pICSL4723OD vector for CRISPR-Cas9 cloning.

## Author contributions

Conceptualization: MS; Investigation: HL. Data analysis: AF, MM, NK. Funding acquisition: MS, RH, NK. Project administration: MS; Supervision: RH, NK, MS; Writing – original draft: HL, MS; Writing – review & editing: all authors.

## Data availability

The raw RNAseq reads are available under the accession number PRJNA1481920 (BioSample accession SAMN61095570 - SAMN61095646) and is publicly accessible using the following link: http://www.ncbi.nlm.nih.gov/bioproject/1481920.Gene counts, TPM values and all results of the DESeq2 analysis are shown in supplementary data table 8. Additionally, all scripts and additional raw data is stored in an ARC structured git, accessible via: https://gitlab.plantmicrobe.de/he_le/Leicher_2026_RNASeq.

## Supplementary figures and legends

**Figure S1:**
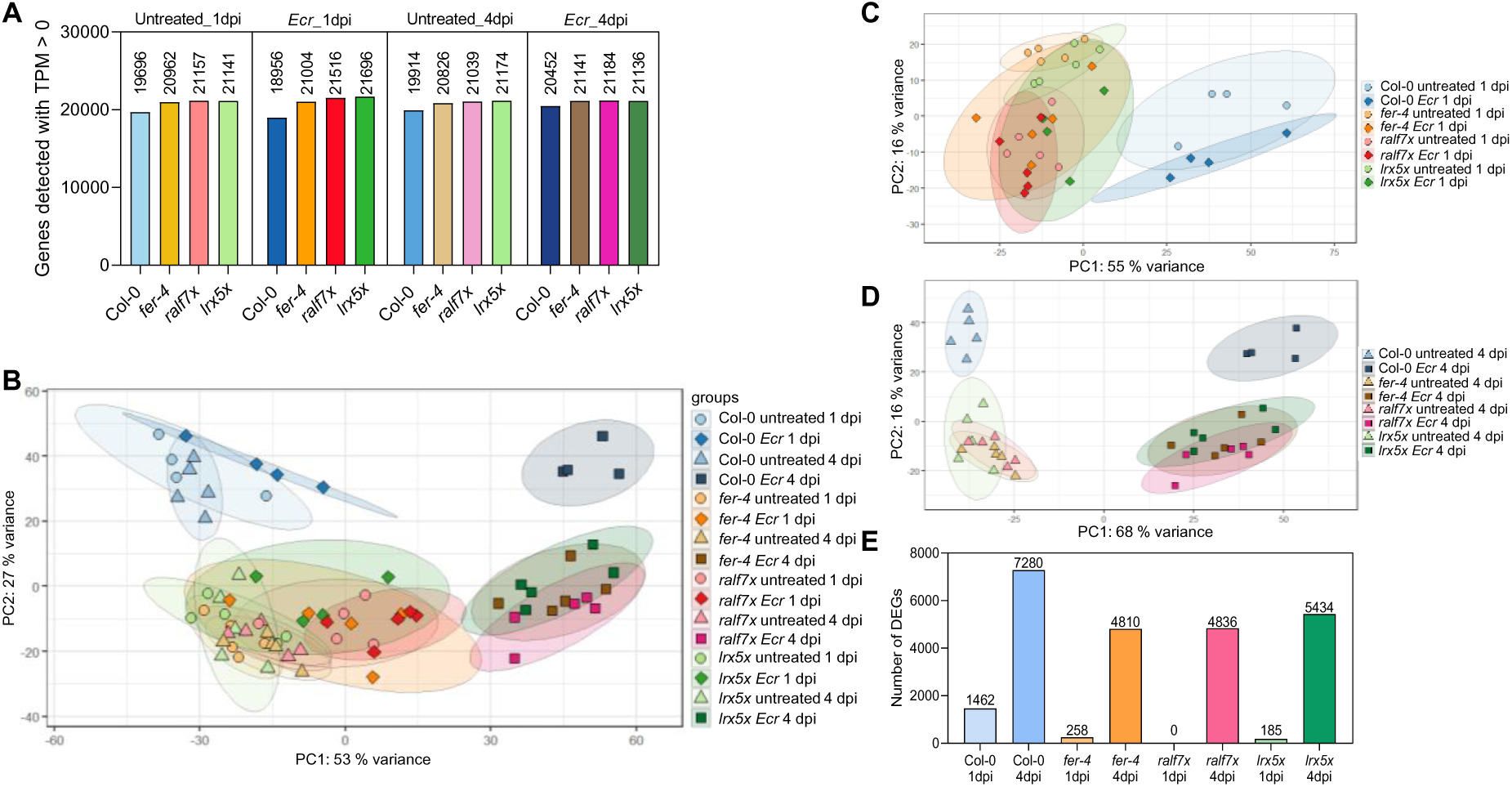
*Ecr* infection leads to transcriptional changes at 1 and 4 dpi. A) Number of detected genes in all tested samples. B) PCA-Analysis of all samples based on the variance stabilized counts from DESeq2. C) PCA-Analysis of samples at 1 dpi based on the variance stabilized counts from DESeq2. D) PCA-Analysis of samples at 4 dpi based on the variance stabilized counts from DESeq2. E) Number of DEGs in all samples (FDR < 0.05, log2(Fold change) > 1 or log2(Fold change) < -1).

**Figure S2:**
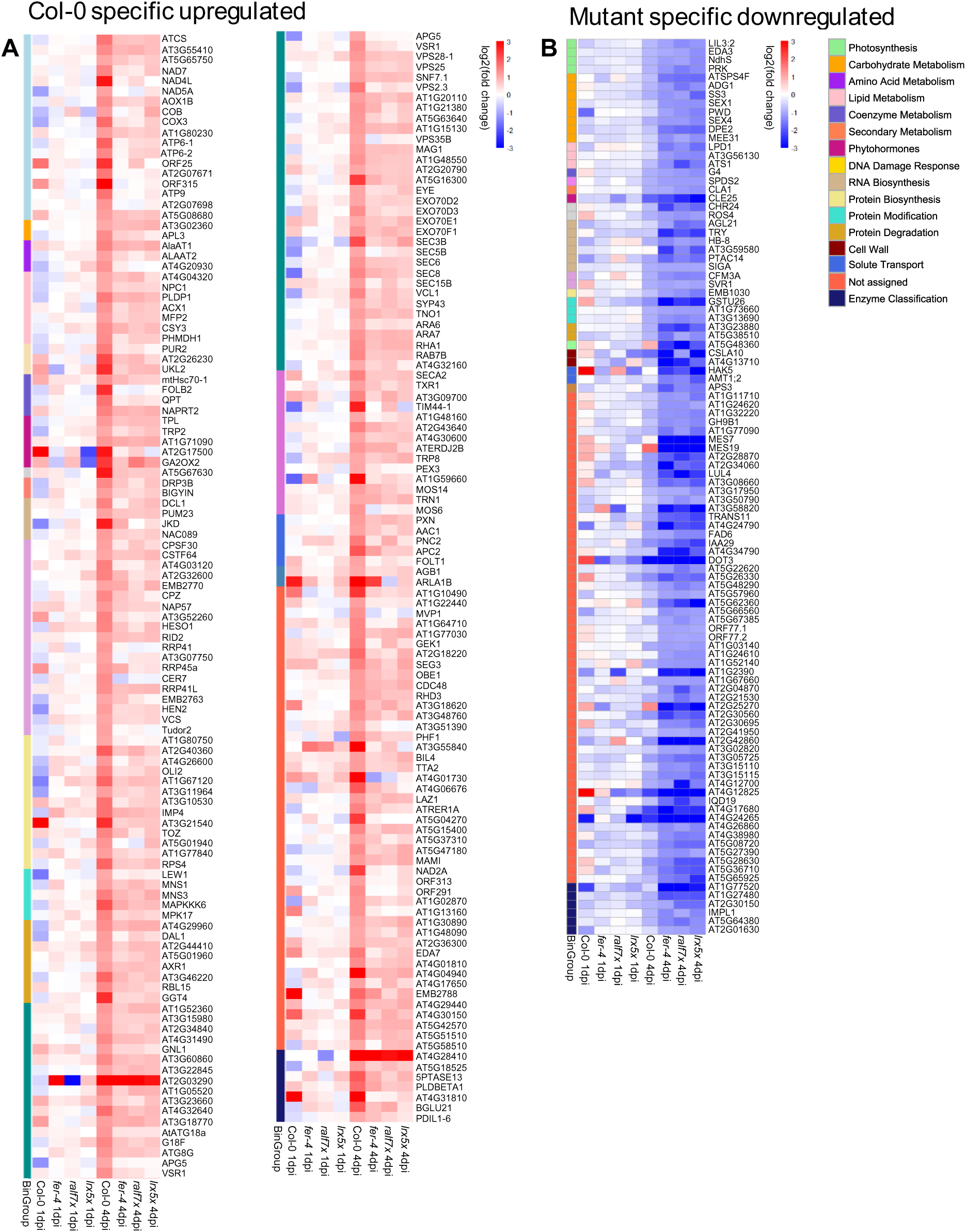
List of Col-0 upregulated and mutants specific downregulated DEGs at 4 dpi. A) Log2(Fold change) of DEGs that are specifically upregulated in Col-0 at 4 dpi but not in the mutant genotypes. B)

**Figure S3:**
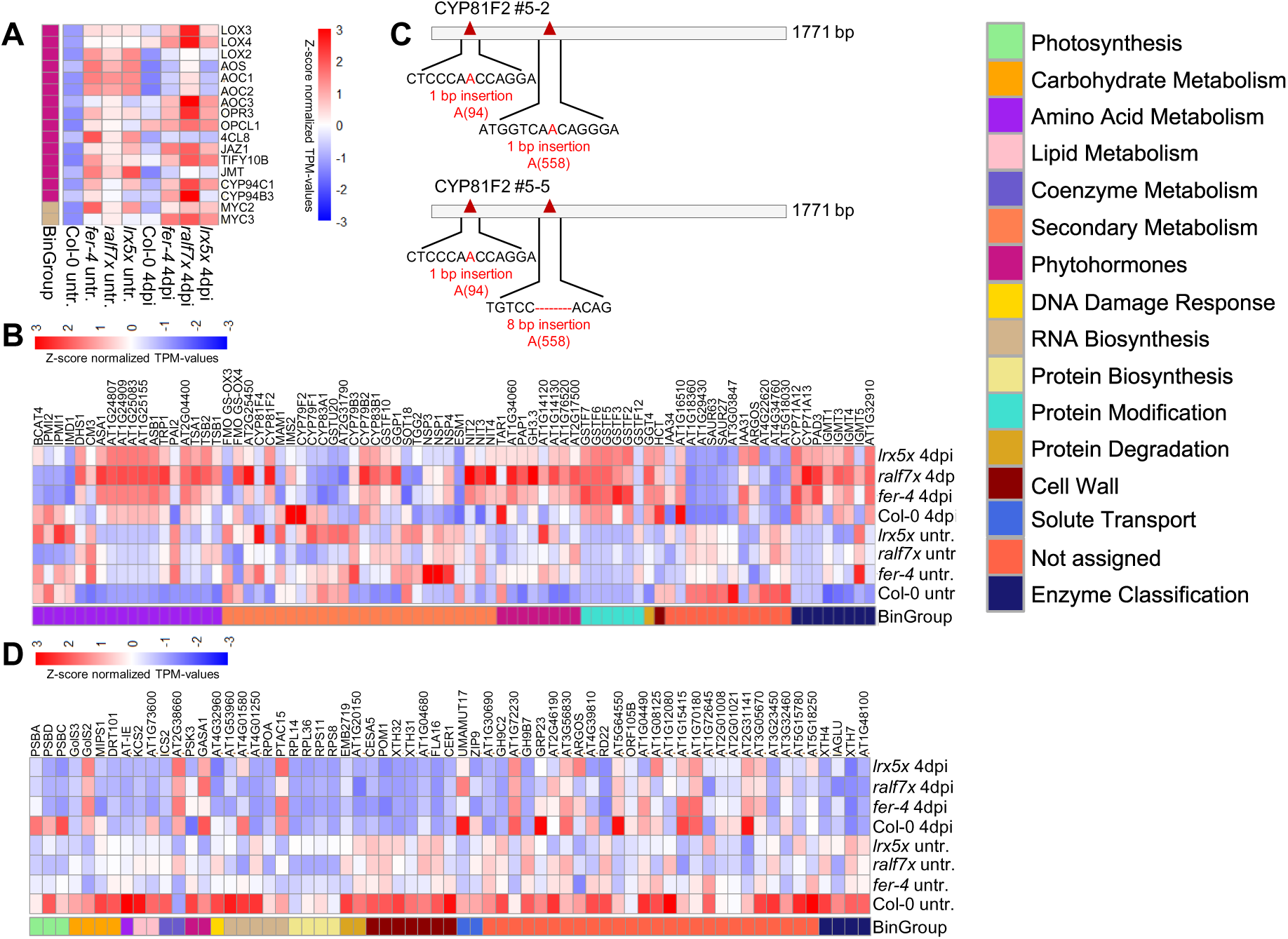
Differences in gene expression levels of genes involved in JA signalling and tryptophane biosynthesis and biosynthesis of tryptophane-derived secondary metabolites. A) Z-score normalized TPM-values of genes involved in JA biosynthesis and signalling. B) Z-score normalized TPM-values of genes involved in the shikimate pathway and biosynthesis of tryptophane derived secondary metabolites. C) Characterization of *fer-4*/CRISPR *cyp81F2* #5-2 and #5-5. Schematic diagram of *CYP81F2* gene structure and the CRISPR-Cas9-mediated mutation pattern detected by DNA sequencing. D) Z-score normalized TPM-values of genes that are significantly less expressed in all mutants compared to Col-0 at the untreated stage.

**Figure S4:**
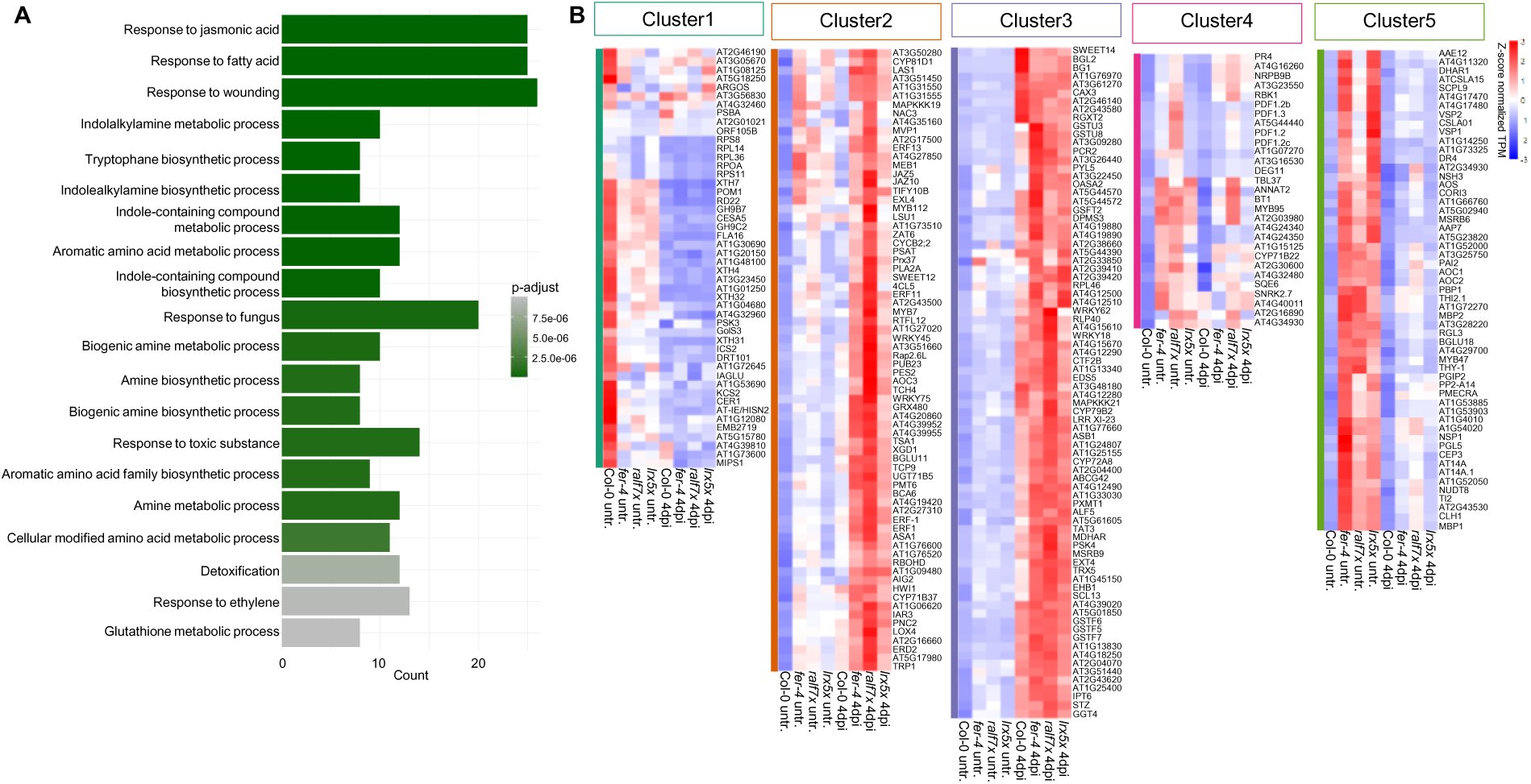
Analysis of the 284 genes that are significantly regulated in Col-0 at 4 dpi and differentially expressed in all mutant genotypes at an untreated stage compared to Col-0. A) Z-score normalized TPM-values of genes that are regulated in Col-0 at 4 dpi and differentially expressed in all mutant genotypes at an untreated stage compared to Col-0. DEGs are grouped in clusters based on Figure 8 B. B) GO-Term analysis of the 284 genes that are regulated in Col-0 at 4 dpi and differentially expressed in all mutant genotypes at an untreated stage compared to Col-0.

## References

Andrews S 2010. FastQC: A Quality Control Tool for High Throughput Sequence Data. http://www.bioinformatics.babraham.ac.uk/projects/fastqc/: Babraham Bioinformatics, Babraham Institute, Cambridge, UK.

Bacete L, Schulz J, Engelsdorf T, Bartosova Z, Vaahtera L, Yan G, Gerhold JM, Tichá T, Øvstebø C, Gigli-Bisceglia N, et al. 2022. THESEUS1 modulates cell wall stiffness and abscisic acid production in Arabidopsis thaliana. Proceedings of the National Academy of Sciences 119: e2119258119.

Bednarek P, Pislewska-Bednarek M, Svatos A, Schneider B, Doubsky J, Mansurova M, Humphry M, Consonni C, Panstruga R, Sanchez-Vallet A, et al. 2009. A glucosinolate metabolism pathway in living plant cells mediates broad-spectrum antifungal defense. Science 323: 101–106.

Berger S, Sinha AK, Roitsch T. 2007. Plant physiology meets phytopathology: plant primary metabolism and plant–pathogen interactions. Journal of Experimental Botany 58: 4019–4026.

Biermann D, von Arx M, Munzert-Eberlein KS, Xhelilaj K, Séré D, Stegmann M, Vert G, Wolf S, Engelsdorf T, Zipfel C, et al. 2025. A RALF-brassinosteroid signaling circuit regulates Arabidopsis hypocotyl cell shape. Current Biology 35: 5002–5017.e5005.

Bolger AM, Lohse M, Usadel B. 2014. Trimmomatic: a flexible trimmer for Illumina sequence data. Bioinformatics 30: 2114–2120.

Cao J, Li M, Chen J, Liu P, Li Z. 2016. Effects of MeJA on Arabidopsis metabolome under endogenous JA deficiency. Scientific Reports 6: 37674.

Carlson M 2024. org.At.tair.db: Genome wide annotation for Arabidopsis_. R package version 3.20.0.

Castel B, Tomlinson L, Locci F, Yang Y, Jones JDG. 2019. Optimization of T-DNA architecture for Cas9-mediated mutagenesis in Arabidopsis. PLOS ONE 14: e0204778.

Chandran D, Inada N, Hather G, Kleindt CK, Wildermuth MC. 2010. Laser microdissection of Arabidopsis cells at the powdery mildew infection site reveals site-specific processes and regulators. Proceedings of the National Academy of Sciences 107: 460–465.

Chen J, Yu F, Xu F. 2024. Not just signals: RALFs as cell wall-structuring peptides. Trends in Plant Science 29: 727–729.

Cheng C-Y, Krishnakumar V, Chan AP, Thibaud-Nissen F, Schobel S, Town CD. 2017. Araport11: a complete reannotation of the Arabidopsis thaliana reference genome. The Plant Journal 89: 789–804.

Cheung AY. 2024. FERONIA: A Receptor Kinase at the Core of a Global Signaling Network. Annual Review of Plant Biology 75: 345–375.

Chiniquy D, Underwood W, Corwin J, Ryan A, Szemenyei H, Lim CC, Stonebloom SH, Birdseye DS, Vogel J, Kliebenstein D, et al. 2019. PMR5, an acetylation protein at the intersection of pectin biosynthesis and defense against fungal pathogens. The Plant Journal 100: 1022–1035.

Chowdhury J, Henderson M, Schweizer P, Burton RA, Fincher GB, Little A. 2014. Differential accumulation of callose, arabinoxylan and cellulose in nonpenetrated versus penetrated papillae on leaves of barley infected with Blumeria graminis f. sp. hordei. New Phytologist 204: 650–660.

Chowdhury J, Lück S, Rajaraman J, Douchkov D, Shirley NJ, Schwerdt JG, Schweizer P, Fincher GB, Burton RA, Little A. 2017. Altered Expression of Genes Implicated in Xylan Biosynthesis Affects Penetration Resistance against Powdery Mildew. Frontiers in Plant Science 8–2017.

Consonni C, Bednarek P, Humphry M, Francocci F, Ferrari S, Harzen A, Ver Loren van Themaat E, Panstruga R. 2009. Tryptophan-Derived Metabolites Are Required for Antifungal Defense in the Arabidopsis mlo2 Mutant. Plant Physiology 152: 1544–1561.

Cosgrove DJ. 2024. Structure and growth of plant cell walls. Nature Reviews Molecular Cell Biology 25: 340–358.

Delgado-Cerezo M, Sánchez-Rodríguez C, Escudero V, Miedes E, Fernández PV, Jordá L, Hernández-Blanco C, Sánchez-Vallet A, Bednarek P, Schulze-Lefert P, et al. 2012. Arabidopsis Heterotrimeric G-protein Regulates Cell Wall Defense and Resistance to Necrotrophic Fungi. Molecular Plant 5: 98–114.

Duan Q, Kita D, Li C, Cheung AY, Wu H-M. 2010. FERONIA receptor-like kinase regulates RHO GTPase signaling of root hair development. Proceedings of the National Academy of Sciences 107: 17821–17826.

Eichmann R, Hückelhoven R. 2008. Accommodation of powdery mildew fungi in intact plant cells. Journal of Plant Physiology 165: 5–18.

Ellis C, Turner JG. 2001. The Arabidopsis Mutant cev1 Has Constitutively Active Jasmonate and Ethylene Signal Pathways and Enhanced Resistance to Pathogens. The Plant Cell 13: 1025–1033.

Endler A, Persson S. 2011. Cellulose Synthases and Synthesis in Arabidopsis. Molecular Plant 4: 199–211.

Engelsdorf T, Horst RJ, Pröls R, Pröschel M, Dietz F, Hückelhoven R, Voll LM. 2013. Reduced Carbohydrate Availability Enhances the Susceptibility of Arabidopsis toward Colletotrichum higginsianum. Plant Physiology 162: 225–238.

Fotopoulos V, Gilbert MJ, Pittman JK, Marvier AC, Buchanan AJ, Sauer N, Hall JL, Williams LE. 2003. The monosaccharide transporter gene, AtSTP4, and the cell-wall invertase, Atbetafruct1, are induced in Arabidopsis during infection with the fungal biotroph Erysiphe cichoracearum. Plant Physiology 132: 821–829.

Gao H, Niu J, Yang X, He D, Wang C. 2014. Impacts of powdery mildew on wheat grain sugar metabolism and starch accumulation in developing grains. Starch - Stärke 66: 947–958.

Glawe DA. 2008. The powdery mildews: a review of the world’s most familiar (yet poorly known) plant pathogens. Annual Review of Phytopathology 46: 27–51.

Guérin A, Levasseur C, Herger A, Renggli D, Sotiropoulos AG, Kadler G, Hou X, Schaufelberger M, Meyer C, Wicker T, et al. 2024. Histidine limitation alters plant development and influences the TOR network. Journal of Experimental Botany 76: 1085–1098.

Guo H, Nolan TM, Song G, Liu S, Xie Z, Chen J, Schnable PS, Walley JW, Yin Y. 2018. FERONIA Receptor Kinase Contributes to Plant Immunity by Suppressing Jasmonic Acid Signaling in Arabidopsis thaliana. Current Biology 28: 3316–3324.e3316.

Herger A, Gupta S, Kadler G, Franck CM, Boisson-Dernier A, Ringli C. 2020. Overlapping functions and protein-protein interactions of LRR-extensins in Arabidopsis. PLOS Genetics 16: e1008847.

Hückelhoven R, Panstruga R. 2011. Cell biology of the plant–powdery mildew interaction. Current Opinion in Plant Biology 14: 738–746.

Hunziker P, Ghareeb H, Wagenknecht L, Crocoll C, Halkier BA, Lipka V, Schulz A. 2020. De novo indol-3-ylmethyl glucosinolate biosynthesis, and not long-distance transport, contributes to defence of Arabidopsis against powdery mildew. Plant, Cell & Environment 43: 1571–1583.

Ingle RA. 2011. Histidine biosynthesis. Arabidopsis Book 9: e0141.

Ishida K, Yokoyama R. 2022. Reconsidering the function of the xyloglucan endotransglucosylase/hydrolase family. Journal of Plant Research 135: 145–156.

Jiang W, Wang Z, Li Y, Liu X, Ren Y, Li C, Luo S, Singh RM, Li Y, Kim C, et al. 2024. FERONIA regulates salt tolerance in Arabidopsis by controlling photorespiratory flux. The Plant Cell 36: 4732–4751.

Johnson LYD, Major IT, Chen Y, Yang C, Vanegas-Cano LJ, Howe GA. 2023. Diversification of JAZ-MYC signaling function in immune metabolism. New Phytologist 239: 2277–2291.

Kai K, Takahashi H, Saga H, Ogawa T, Kanaya S, Ohta D. 2011. Metabolomic characterization of the possible involvement of a Cytochrome P450, CYP81F4, in the biosynthesis of indolic glucosinolate in Arabidopsis. Plant Biotechnology 28: 379–385.

Kessler SA, Shimosato-Asano H, Keinath NF, Wuest SE, Ingram G, Panstruga R, Grossniklaus U. 2010. Conserved molecular components for pollen tube reception and fungal invasion. Science 330: 968–971.

Kim D, Paggi JM, Park C, Bennett C, Salzberg SL. 2019. Graph-based genome alignment and genotyping with HISAT2 and HISAT-genotype. Nature Biotechnology 37: 907–915.

Kolde R 2025. pheatmap: Pretty Heatmaps. https://CRAN.R-project.org/package=pheatmap.

Kusch S, Panstruga R. 2017. mlo-Based Resistance: An Apparently Universal “Weapon” to Defeat Powdery Mildew Disease. Molecular Plant-Microbe Interactions 30: 179–189.

Lamesch P, Berardini TZ, Li D, Swarbreck D, Wilks C, Sasidharan R, Muller R, Dreher K, Alexander DL, Garcia-Hernandez M, et al. 2012. The Arabidopsis Information Resource (TAIR): improved gene annotation and new tools. Nucleic Acids Research 40: D1202–D1210.

Leicher H, Schade SD, Huebbers JW, Munzert-Eberlein KS, Haljiti G, Biermann D, Makris A, Zhu X, Chauhan Y, Ludwig C, et al. 2026. Endogenous RALF peptide function is required for powdery mildew host colonization. New Phytologist in press.

Li H, Handsaker B, Wysoker A, Fennell T, Ruan J, Homer N, Marth G, Abecasis G, Durbin R, Subgroup GPDP. 2009. The Sequence Alignment/Map format and SAMtools. Bioinformatics 25: 2078–2079.

Li L, Chen H, Alotaibi SS, Pěnčík A, Adamowski M, Novák O, Friml J. 2022. RALF1 peptide triggers biphasic root growth inhibition upstream of auxin biosynthesis. Proceedings of the National Academy of Sciences 119: e2121058119.

Li Y, Guo G, Zhou L, Chen Y, Zong Y, Huang J, Lu R, Liu C. 2020. Transcriptome Analysis Identifies Candidate Genes and Functional Pathways Controlling the Response of Two Contrasting Barley Varieties to Powdery Mildew Infection. International Journal of Molecular Sciences 21: 151.

Liao Y, Smyth GK, Shi W. 2014. featureCounts: an efficient general purpose program for assigning sequence reads to genomic features. Bioinformatics 30: 923–930.

Lipka U, Fuchs R, Lipka V. 2008. Arabidopsis non-host resistance to powdery mildews. Current Opinion in Plant Biology 11: 404–411.

Liu S, Bartnikas LM, Volko SM, Ausubel FM, Tang D. 2016. Mutation of the Glucosinolate Biosynthesis Enzyme Cytochrome P450 83A1 Monooxygenase Increases Camalexin Accumulation and Powdery Mildew Resistance. Frontiers in Plant Science 7–2016.

Lopa FR, Snigdha FN, Lumactud RA, Sikder MM. 2025. Harnessing Camalexin as a Sustainable and Ecofriendly Strategy to Control Harmful Phytopathogens. Plant Pathology 74: 2463–2477.

Love MI, Huber W, Anders S. 2014. Moderated estimation of fold change and dispersion for RNA-seq data with DESeq2. Genome Biology 15: 550.

Micali C, Göllner K, Humphry M, Consonni C, Panstruga R. 2008. The Powdery Mildew Disease of Arabidopsis: A Paradigm for the Interaction between Plants and Biotrophic Fungi. Arabidopsis Book 6: e0115.

Mielke S, Gasperini D. 2019. Interplay between Plant Cell Walls and Jasmonate Production. Plant and Cell Physiology 60: 2629–2637.

Moussu S, Lee HK, Haas KT, Broyart C, Rathgeb U, De Bellis D, Levasseur T, Schoenaers S, Fernandez GS, Grossniklaus U, et al. 2023. Plant cell wall patterning and expansion mediated by protein-peptide-polysaccharide interaction. Science 382: 719–725.

Muller E, Ropitaux M, Durambur G, Laplaud V, Gebhard L, Lehner A, Drevensek S, Boudaoud A. 2025. Dual role of the receptor kinase FERONIA in regulating tissue mechanics and growth. bioRxiv: 2025.2008.2028.672783.

Proels RK, Hückelhoven R. 2014. Cell-wall invertases, key enzymes in the modulation of plant metabolism during defence responses. Molecular Plant Pathology 15: 858–864.

Roitsch T, González M-C. 2004. Function and regulation of plant invertases: sweet sensations. Trends in Plant Science 9: 606–613.

Rose JK, Braam J, Fry SC, Nishitani K. 2002. The XTH family of enzymes involved in xyloglucan endotransglucosylation and endohydrolysis: current perspectives and a new unifying nomenclature. Plant and Cell Physiology 43: 1421–1435.

Rößling A-K, Dünser K, Liu C, Lauw S, Rodriguez-Franco M, Kalmbach L, Barbez E, Kleine-Vehn J. 2024. Pectin methylesterase activity is required for RALF1 peptide signalling output. Elife 13: RP96943.

Sánchez-Rodríguez C, Bauer S, Hématy K, Saxe F, Ibáñez AB, Vodermaier V, Konlechner C, Sampathkumar A, Rüggeberg M, Aichinger E, et al. 2012. CHITINASE-LIKE1/POM-POM1 and Its Homolog CTL2 Are Glucan-Interacting Proteins Important for Cellulose Biosynthesis in Arabidopsis. The Plant Cell 24: 589–607.

Schade S, Leicher H, von Arx M, Monte I, Gronnier J, Stegmann M. 2025. The interplay of RALF structural and signaling functions in plant-microbe interactions. PLOS Pathogens 21: e1013588.

Schoenaers S, Lee HK, Gonneau M, Faucher E, Levasseur T, Akary E, Claeijs N, Moussu S, Broyart C, Balcerowicz D. 2024. Rapid alkalinization factor 22 has a structural and signalling role in root hair cell wall assembly. Nature Plants 10: 494–511.

Schwacke R, Ponce-Soto GY, Krause K, Bolger AM, Arsova B, Hallab A, Gruden K, Stitt M, Bolger ME, Usadel B. 2019. MapMan4: A Refined Protein Classification and Annotation Framework Applicable to Multi-Omics Data Analysis. Molecular Plant 12: 879–892.

Song L, Xu G, Li T, Zhou H, Lin Q, Chen J, Wang L, Wu D, Li X, Wang L, et al. 2022. The RALF1-FERONIA complex interacts with and activates TOR signaling in response to low nutrients. Molecular Plant 15: 1120–1136.

Stegmann M, Monaghan J, Smakowska-Luzan E, Rovenich H, Lehner A, Holton N, Belkhadir Y, Zipfel C. 2017. The receptor kinase FER is a RALF-regulated scaffold controlling plant immune signaling. Science 355: 287–289.

Sutton PN, Gilbert MJ, Williams LE, Hall JL. 2007. Powdery mildew infection of wheat leaves changes host solute transport and invertase activity. Physiologia Plantarum 129: 787–795.

Swarbrick PJ, Schulze-Lefert P, Scholes JD. 2006. Metabolic consequences of susceptibility and resistance (race-specific and broad-spectrum) in barley leaves challenged with powdery mildew. *Plant*, Cell & Environment 29: 1061–1076.

Vogel JP, Raab TK, Schiff C, Somerville SC. 2002. PMR6, a Pectate Lyase–Like Gene Required for Powdery Mildew Susceptibility in Arabidopsis. The Plant Cell 14: 2095–2106.

Vogel JP, Raab TK, Somerville CR, Somerville SC. 2004. Mutations in PMR5 result in powdery mildew resistance and altered cell wall composition. The Plant Journal 40: 968–978.

Wang P, Clark NM, Nolan TM, Song G, Bartz PM, Liao CY, Montes-Serey C, Katz E, Polko JK, Kieber JJ, et al. 2022. Integrated omics reveal novel functions and underlying mechanisms of the receptor kinase FERONIA in Arabidopsis thaliana. The Plant Cell 34: 2594–2614.

Wright DP, Baldwin BC, Shephard MC, Scholes JD. 1995. Source-sink relationships in wheat leaves infected with powdery mildew. I. Alterations in carbohydrate metabolism. Physiological and Molecular Plant Pathology 47: 237–253.

Wu Y, Sexton WK, Zhang Q, Bloodgood D, Wu Y, Hooks C, Coker F, Vasquez A, Wei C-I, Xiao S. 2023. Leaf abaxial immunity to powdery mildew in Arabidopsis is conferred by multiple defense mechanisms. Journal of Experimental Botany 75: 1465–1478.

Yang T, Wang L, Li C, Liu Y, Zhu S, Qi Y, Liu X, Lin Q, Luan S, Yu F. 2015. Receptor protein kinase FERONIA controls leaf starch accumulation by interacting with glyceraldehyde-3-phosphate dehydrogenase. Biochemical and Biophysical Research Communications 465: 77–82.

Yeats TH, Sorek H, Wemmer DE, Somerville CR. 2016. Cellulose Deficiency Is Enhanced on Hyper Accumulation of Sucrose by a H+-Coupled Sucrose Symporter. Plant Physiology 171: 110–124.

Yu G. 2024. Thirteen years of clusterProfiler. The Innovation 5: 100722.

Yu G, Zhang L, Xue H, Chen Y, Liu X, Del Pozo JC, Zhao C, Lozano-Duran R, Macho AP. 2024. Cell wall-mediated root development is targeted by a soil-borne bacterial pathogen to promote infection. Cell Reports 43: 114179.

Zander M, Lewsey MG, Clark NM, Yin L, Bartlett A, Saldierna Guzmán JP, Hann E, Langford AE, Jow B, Wise A, et al. 2020. Integrated multi-omics framework of the plant response to jasmonic acid. Nature Plants 6: 290–302.

Zeng F-S, Menardo F, Xue M-F, Zhang X-J, Gong S-J, Yang L-J, Shi W-Q, Yu D-Z. 2017. Transcriptome Analyses Shed New Insights into Primary Metabolism and Regulation of Blumeria graminis f. sp. tritici during Conidiation. Frontiers in Plant Science Volume 8 - 2017.

Zhang X, Yang Z, Wu D, Yu F. 2020. RALF-FERONIA Signaling: Linking Plant Immune Response with Cell Growth. Plant Communications 1: 100084.

